# Precision Fermentation Gets a New Player: Genome Assembly and Genetic Transformation of a Novel Strain, *Pichia hangzhouana*, for Precision Fermentation of Food Ingredients

**DOI:** 10.64898/2026.11.19.767948

**Authors:** Ema Johnston, Parastoo Karimian, Ulduz Vafadarshamasbi, Roya Afshari, Andreas Bachler, Oliver Mead, Pavel Boer, Phil Hands, Mya Myintzu Hlaing, Catherine Mary McAuley, Annette Kasprzak, Jian-Wei Liu, Thomas Vanhercke, Carol Janet Hartley

**Affiliations:** CSIRO Agriculture and Food, GPO Box 1700, Acton, ACT 2601, Australia; CSIRO Agriculture and Food, Werribee, Vic, 3030, Australia; CSIRO Environment, GPO Box 1700, Acton, ACT 2601, Australia; School of Agriculture and Food Sustainability, The University of Queensland, St Lucia, QLD 4072, Australia

**Keywords:** yeast, precision fermentation, food ingredients, *Pichia hangzhouana*, *Zygoascus hellenicus*, phylogenomics

## Abstract

Precision fermentation leverages the metabolic pathways of microorganisms to produce valuable proteins and enzymes. *Pichia hangzhouana*, synonymous with *Zygoascus hellenicus*, is a non-conventional yeast with potential for food biotechnological applications. This study characterizes the CSIRO strain *P. hangzhouana FRR5246*, isolated from a tea beverage, to evaluate its suitability as a precision fermentation host for a variety of industrial, agricultural and food production applications.

*P. hangzhouana* demonstrated robust growth across a range of carbon sources, with the highest optical density observed in media containing sucrose or sorbitol as sole carbon source. The yeast showed susceptibility to geneticin, hygromycin B, nourseothricin sulfate, and zeocin sulfate antibiotic agents. Successful genetic transformation was achieved, with the expression of a recombinant orange fluorescent protein (OFP) under the control of the native glyceraldehyde-3-phosphate dehydrogenase (GAP1) promoter. The secretome exhibited strong antimicrobial activity against *Escherichia coli* and *Staphylococcus aureus*, indicating the presence of potent antimicrobial compounds.

In addition, whole genome sequencing of *P. hangzhouana FRR5246* provided genetic tools for recombinant expression of heterologous genes, insight into the genes, genetic architecture and regulatory mechanisms underlying phenotypic qualities, as well as confirming phylogenetic classification of this strain as clustering with the *Dipodascomycetes,* most closely related to species *Zygoascus hellenicus*.

*P. hangzhouana FRR5246* is a promising candidate for precision fermentation of food ingredients, capable of utilizing diverse carbon sources and producing recombinant proteins. In addition, production of antimicrobial compounds further enhances the potential for biotechnological exploitation. Future work will focus on optimizing growth conditions and exploring native inducible promoters for controlled gene expression.

## 1. Introduction

Precision fermentation utilizes native and edited metabolic pathways of micro-organisms to produce natural and recombinant protein and protein products useful for industrial applications in agriculture, food and the pharmaceutical industry(1, 2) . At the forefront of this growing area of research is the development of non-conventional fungal and bacterial strains that have the potential to produce native and recombinant proteins or enzymes at scale. The first step in this process is to characterize these non-conventional strains and determine if they are suitable hosts as expression systems for industrial applications(3). Understanding the genetics, carbon utilization and genetic malleability of these strains is essential for this process (4).

*Pichia hangzhouana* (*P. hangzhouana*), synonymous with *Zygoascus hellenicus,* remains virtually untouched by synthetic biology. *P. hangzhouana* has been isolated from a range of environments, including tea beverages, spoiled and rotten grapes, fermented food, and human blood cultures (5–8). This yeast has potential for biotechnological applications, given that it has the ability to grow in vitamin-free media across a range of pH levels and can assimilate a wide variety of carbon and nitrogen sources, including a number of agrifood-waste streams (7). To date, *P. hangzhouana* has been used to produce and secrete exo-polygalacturonase when grown on orange peel waste, demonstrating the potential biotechnological importance of *P. hangzhouana* (3).

The strain characterized in this paper, *P. hangzhouana* CSIRO5246, was isolated from a tea beverage in 1998, and was collected and housed in the CSIRO Food Microbiology (FRR) culture collection (9).

Phylogenetic analyses and genetic similarity have shown a close relationship between *Pichia hangzhouana* and *Zygoascus meyerae*, suggesting a possible evolutionary link or functional resemblance between the two species (10). In this work we have assembled a high-quality genome to enable the identification of key genetic features that contribute to the metabolic flexibility of *P. hangzhouana* and the discovery of novel genes and pathways that can be harnessed for biotechnological purposes.

We further characterize the CSIRO yeast strain *P. hangzhouana* FRR5246 when cultured using a variety of alternate carbon sources and assessed susceptibility to a range of antibiotics. Literature has already documented that *P. hangzhouana* has the ability to use a variety of different nitrogen sources and it may be advantageous to combine this ability with a potential waste stream source of carbon to produce cellular biomass, as well as recombinant and non-recombinant proteins of interest to the food and agriculture industries (3). In addition, we further demonstrate herein that the concentrated secretome from *P. hangzhouana* contains antimicrobial activity. Finally, we demonstrate the genetic transformation of *P. hangzhouana* and expression of a recombinant fluorescent protein (OFP).

## 2. Methods and Materials

### Strains and growth conditions

The strain used in this study was the *P. hangzhouana FRR5246* strain from the CSIRO FRR culture collection (CSIRO, Australia). NEB 5 alpha competent *E. coli* cells (NEB, Australia) were used for cloning. Cultivations were conducted in Luria Broth (LB) media for *E. coli* and yeast cultures were either grown in YPD medium (1% w/v yeast extract, 2% w/v peptone, and 2% w/v glucose), buffered minimal dextrose (BMD) medium (1.34% Yeast Nitrogen Base (YNB), 4 × 10⁻⁵% biotin, 200 mM potassium phosphate buffer pH 6.0, and 2% glucose), and buffered minimal methanol (BMM) medium (1.34% YNB, 4 × 10⁻⁵% biotin, 200 mM potassium phosphate buffer pH 6.0), with the addition of 2% or as specified in the manuscript of the chosen carbon source. Antibiotic Kanamycin (Invitrogen) was added to the media when required at a final concentration of 50 µg/mL for *E. coli*. Antibiotics Geneticin (GoldBio), Nourseothricin (GoldBio), Zeocin (Invitrogen), and Hygromycin B (Invitrogen) were added to the media when required at a final concentration of 400 µg/mL, 100 µg/mL, 100µg/mL, and 250µg/mL, respectively. YPD plate cultures were grown in a 30°C stationary incubator.

### Imaging light microscopy and SEM

For Scanning Electron Microscopy (SEM) 1 ml of culture was pelleted via centrifugation at 2000 rpm for 5 minutes and growth media removed. Cells were fixed in formalin–acetic acid–alcohol (FAA; 3.7% formaldehyde v/v, 5% acetic acid, 50% ethanol) overnight, dehydrated through 70%, 80%, 90% and 100% ethanol series, each for 15 mins, before critical point drying in an Autosamdri-815 automatic critical point drier (Tousimis Research Corporation, Rockville USA). Samples were mounted onto double-sided adhesive carbon tabs (ProSciTech, Townsville City, QLD, Australia) and excess, unadhered sample removed using a jet of compressed air. Samples were gold coated using 0a MiniQS sputter coater (Quorum Technologies Ltd., EastSussex, UK) for 30 seconds under an argon environment and imaged using a Zeiss EVO LS 15 scanning electron microscope (Carl Zeiss Microscopy GmbH, Jena, Germany) under high vacuum using a secondary electron detector with 27 kV accelerating voltage.

Brightfield and fluorescence images were obtained using a Zeiss AxioZoom V.16 microscope equipped with white light and fluorescence illumination (HXP 200 C), a PlanNeoFluar 1.0x NA = 0.25 objective and Zeiss Axiocam 712 colour CCD camera (Carl Zeiss Micro-imaging GmbH, Jena, Germany). Orange fluorescent Protein expression (OFP) in transformed *P. hangzhouana* colonies was detected using Zeiss filter set 43 HE (excitation BP550/25; beam splitter FT570; emission BP 605/70). Extended depth of focus image capture and processing was carried out using ZEN blue v3.5 (Carl Zeiss Micro-imaging GmbH, Jena, Germany).

### Organism classification verification using nuclear ribosomal DNA ITS1 to ITS4 region

Axenic cultures of *P. hangzhouana* were grown on YPD medium for 3 days at 30°C. Biomass was collected by scraping approximately 100 mg of cells into a 2 mL tube and genomic DNA extracted using the Quick-DNA Fungal/Bacterial Miniprep Kit (Zymo Research, USA). DNA gel electrophoresis verified the quality of the chromosomal DNA, providing a clean, unfragmented, high-molecular weight band. Nanodrop spectroscopy indicated the purity and concentration of DNA.

To confirm the identification of the *P. hangzhouana* FRR5246 strain, the ITS1--ITS4 region of the nuclear ribosomal DNA (rDNA) was amplified via polymerase chain reactions (PCR) using universal primers ITS1 (5′TCCGTAGGTGAACCTGCGG) and ITS4 (5′TCCTCCGCTTATTGATATGC) (Raja *et al.* <u>2017</u>). The PCR was conducted with OneTaq® 2X Master Mix with Standard Buffer (New England Biolabs, USA), 6µL of genomic DNA was added, along with 1µL of each of the forward and reverse primers at a concentration of 10 µM each, and UltraPure™ Dnase/Rnase-Free Distilled Water up to a final PCR volume of 50 µL (ThermoFisher, 10,977,015). The PCR reaction was prepared by following OneTaq® 2X Master Mix NEB protocols. The PCR began with an initial denaturation step at 94 °C for 30 seconds, followed by 35 cycles of 94 °C for 30 seconds, then 55°C for 30 seconds, the template extension step at 68°C for 1 minute, and a final 5-minute extension cycle at 68 °C. The PCR was carried out on a Kiratec PCR Super Cycler instrument.

The purification used 50 µL of the PCR product and followed the manufacturer’s instructions using an ISOLATE II PCR and Gel Kit (Meridian Biosciences, USA). The purified PCR product was diluted with 30 µL of UltraPure™ Dnase/Rnase-Free Distilled Water (ThermoFisher, 10,977,015) and stored at −20 °C. For Sanger sequencing, PCR amplicons were sent to the Biomolecular Resource Facility at John Curtin School of Medical Research, Australian National University (ANU), Canberra, Australia.

### Analysis of ITS sequencing

Sequencing data was inspected, trimmed, and quality-checked using Geneious Prime 2022 (Biomatters Ltd., Auckland, New Zealand). Low-quality and ambiguous base calls were removed prior to analysis. Consensus sequences were generated and compared against sequences available in the National Centre for Biotechnology Information (NCBI) nucleotide database using the Basic Local Alignment Search Tool (BLASTn) (https://blast.ncbi.nlm.nih.gov/). Taxonomic identification was based on sequence similarity to reference sequences in the NCBI database.

### Proteome Characterisation by liquid chromatography-mass spectrometry (LC-MS) Proteomics

Proteins present in the culture supernatant were precipitated using 4X volumes of cold acetone and incubated at −20 °C overnight. After precipitation, protein pellet was resuspended in 8M urea containing 20mM ammonium bicarbonate. Protein concentration was determined using a Braford assay kit according to the manufacture (Bio-Rad) using bovine serum albumin (BSA) as a standard.

Approximately 10 µL of the protein solution (equivalent to 5 µg) was reduced with 1.0 µL of 15 % (w/v) dithiothreitol (DTT) for 30 min at room temperature, followed by alkylation with 1.0 µL of 40 % (w/v) acrylamide under the same conditions. Protein digestion was performed by adding 87 µL of sequencing-grade trypsin solution (0.1 µg trypsin in 25 mM ammonium bicarbonate) and incubating at 37 °C for 3 to 4 hours. The reaction was quenched with 1 µL of 10 % (v/v) formic acid, and the digested peptides were clarified using a 0.22 µm filter.

For LC–MS analysis, 100 ng of desalted tryptic peptides was loaded onto a trap column (PepMap100 C18, 5 mm × 300 µm, 5 µm; Thermo Scientific) and subsequently separated on a nano analytical column (PepMap100 C18, 150 mm × 75 µm, 2 µm; Thermo Scientific) using an Ultimate 3000 RSLC nano LC system (Thermo Scientific). Mobile phase A consisted of water with 0.1 % (v/v) formic acid, and mobile phase B contained 80 % (v/v) acetonitrile with 0.08 % (v/v) formic acid. Peptides were eluted with a linear gradient from 5 % to 50 % B over 50 min, followed by an increase to 99 % B over 10 min. Eluted peptides were ionized using a Nanospray Flex Ion Source (Thermo Scientific) operating at 2.3 kV and 300 °C.

Mass spectrometric analysis was carried out on an Orbitrap Fusion MS (Thermo Scientific). Survey scans of precursor ions (MS1) were acquired in the Orbitrap at 120 K resolution (400–1500 m/z), with an AGC target of 4 × 10⁵ and a maximum injection time of 50 ms. The most intense precursors (charge states 2+ to 7+) above 1 × 10⁵ intensity were selected for fragmentation by high-energy collisional dissociation (HCD, 28 % energy). Fragment ions (MS/MS) were detected in the ion trap at a rapid scan rate, with AGC = 4 × 10³, maximum injection time = 300 ms, and a dynamic exclusion period of 15 s. The instrument operated in top-speed mode with a 3 s cycle for MS/MS acquisition.

Protein identification and quantification were performed using Proteome Discoverer 2.2 (Thermo Scientific) with the Sequest HT search engine. Spectral data were searched against the protein sequences of *P. hangzhouana*. The precursor mass tolerance was set to 10 ppm and product ion tolerance to 0.6 Da, allowing up to three missed tryptic cleavages. Variable modifications included oxidation (+15.995 Da), deamidation (+0.984 Da), amidation (−0.984 Da), and propionamidation (+71.037 Da). Peptide-spectrum matches were validated using the Percolator algorithm at a 1 % false discovery rate (FDR). Relative protein abundance was calculated from precursor ion intensities and normalized to total peptide content.

### Secretome analysis

To compare the changes in the secretome between 24 and 48 h, normalised protein abundances were averaged across replicate et each time point. Descriptive log₂ fold changes were then calculated for proteins detected at both 24 h and 48 h as log₂ (48 h/24 h). The ten proteins with the largest absolute log₂ fold-change values were selected for visualisation.

Signal peptides were predicted using SignalP 6.0 (http://www.cbs.dtu.dk/services/SignalP). This tool predicts the presence of signal peptides and their likely cleavage sites in amino acid sequences, enabling classification of proteins with predicted secretion signals.

Secretome proteins were functionally interpreted by retrieving their corresponding annotations from the genome-wide eggNOG-mapper/InterProScan annotation table using gene identifiers. For GO-based functional classification, GO identifiers were extracted from the combined GO annotation field and mapped to GO term names and ontology categories using the GO.db and AnnotationDbi packages in *R*. Duplicate GO terms within each protein were collapsed, and GO term frequency was calculated as the number of proteins associated with each Biological Process or Molecular Function term.

### Antimicrobial testing

#### Preparation of *Pichia* supernatant

*Pichia hangzhouana* was cultivated according to the conditions detailed in Supp Table **x**. Approximately three to five morphologically similar colonies from the agar plates were transferred into 20 mL of YPD broth in 50 mL Falcon centrifuge tubes and incubated for 48 hours at 30 °C with shaking at 250 rpm. The supernatant was collected by centrifugation at 12,000 *g*, filter sterilized using 0.2 μm filters and concentrated with a SpeedVac Vacuum Concentrator (Thermo Fisher). After overnight concentration, the supernatant was resuspended in 2 mL of YPD broth media, and the total crude protein was measured using a NanoDrop 2000c spectrophotometer.

### Preparation of bacterial cells stocks and inoculum

*Escherichia coli (E. coli, ATCC 25922) and Staphylococcus aureus (S. aureus, ATCC 25923)* cultures were prepared using the conditions noted in Supp Table **x**. Three to five morphologically similar colonies from the Nutrient agar plates were suspended in 10 mL Nutrient broth and incubated overnight at 37 °C, statically. The overnight culture was collected by centrifugation, washed two times with sterilised MilliQ water and resuspended in Mueller–Hinton (MH) broth. The cell suspension was adjusted to give a turbidity equivalent to 1 × 10^9^ cfu/mL (where the OD_600_ value=1 was adjusted to the equivalent of 10^9^). The actual number of colony-forming units was confirmed from the viability plate count. Suspensions of 5 × 10^5^ cfu/mL (final concentration) were used for the broth dilutions of antimicrobial testing. *et al*.

### MIC using microtiter assay plate method

*In-vitro* antimicrobial activity against *E. coli*, ATCC 25922, and *S. aureus*, ATCC 25923, were tested by using susceptibility micro dilution assay according to CLSI (Clinical and Laboratory Standards Institute approved guidelines) as described elsewhere (CLSI, 2019) and by minimum inhibitory concentration (MIC) determination for each microorganism of interest. The MIC were determined visually and spectrophotometrically in the broth dilutions as the lowest concentrations of the *P. hangzhouana* supernatant at which no bacterial growth was observed. The antibacterial activity of the *P. hangzhouana* supernatant was studied by employing a microdilution method, using MH broth. The concentrated *P. hangzhouana* supernatant was diluted with culture broth to a stock concentration. The *Pichia* supernatant prepared from CSIRO FRR strain 5246 was tested at concentrations ranging from 180 - 1.4 mg/mL. Ciprofloxacin was used as an antibiotic control at concentrations ranging from 2 µg/mL to 0.0156 µg/mL. Sterilised MH broth was used as the diluent broth for the assay. The stock solutions of *Pichia* supernatant and the ciprofloxacin control were prepared to the desired concentration to allow dilution with a volume of microbial suspensions. Briefly, diluted *Pichia* supernatant suspension (90 µL) was transferred across the row in all the microtitre plate test wells with descending concentrations. Then 10 µL of the ∼ 10^6^ CFU/mL inoculum prepared in the broth was transferred into all test wells to give the desired final inoculum load of ∼10^5^ CFU/mL. Each plate had a set of controls including the antibiotic control (ciprofloxacin, 2-0.0156 µg/mL, in serial dilution), the positive growth control (broth inoculated with *E. coli* or *S. aureus* and no *Pichia* supernatant) and the sterile negative control (the sterile broth and *Pichia* supernatant).

Each plate was wrapped with parafilm to ensure that cultures did not become dehydrated. The plates were prepared in triplicate and incubated using the growth conditions provided in Supp Table **x**. After the incubation, antimicrobial activity was determined by the total biomass of the treatment culture in each well using optical density (OD) measurements at λ= 600 nm with a spectrophotometer microplate reader (Thermo Scientific Varioskan® Flash). The mean absolute change of OD in the *Pichia* supernatant-treated sample compared to the OD of the day 0 sample was calculated for evaluating each microbial growth in each treatment. The lowest concentration at which the OD change occurred was taken as the MIC value. Data from treated samples of each strain were compared with the growth control without antimicrobial agents. The percentage of inhibition (I, %) of the *Pichia* supernatant was calculated according to the formula: I (%) = (OD_C_ − OD_T_/OD_C_) ×100, where OD_C_ is the optical density of the growth control and OD_T_ is the optical density of the tested microbial growth in each well containing *Pichia* supernatant.

### MBC

The minimum bactericidal concentration (MBC) was determined by subculturing the wells (with no OD value change) from the broth dilution of MIC tests onto fresh Nutrient agar plates. After the incubation, the lowest concentration that did not show any visible growth was taken as the MBC.

### MIC using Disk Diffusion method

*In-vitro* antimicrobial activity against *E. coli*, ATCC 25922, and *S. aureus*, ATCC 25923, were tested by Disk Diffusion Susceptibility according to CLSI (Clinical and Laboratory Standards Institute approved guidelines) as described elsewhere (11) (CLSI, 2019) and by MIC (Minimum Inhibitory Concentration) determination for each microorganism of interest. The test is performed by applying a bacterial inoculum of approximately 1–2 × 10^8^ CFU/mL to the MH agar plate (92 mm diameter). Paper antibiotic disks were inoculated with 100 µL of *Pichia* supernatant at a concentration of 180 mg/mL (*P. hangzhouana* strain 5246). The control disks inoculated with 40 µL of 250 µg/mL Ciprofloxacin were used as MIC controls. The disks were placed in the centre of the agar surface inoculated with *E. coli* or *S. aureus*. The plates were incubated at 37 °C for 18–24 hours. After incubation, the zones of bacterial growth inhibition around each disk were measured to the nearest millimetre. The diameter of the inhibition zones reflected the bacterial susceptibility and was influenced by the diffusion rate of the *Pichia* supernatant through the agar. The results were interpreted according to CLSI standards for antimicrobial susceptibility testing.

### Genomic DNA extraction

Axenic cultures of *P. hangzhouana* were grown on YPD medium for 3 days at 30°C. Biomass was collected by scraping approximately 100 mg of cells into a 2 mL tube and genomic DNA extracted using the DNA HMW MagBead Kit (Zymo Research, USA). DNA gel electrophoresis verified the quality of the chromosomal DNA, providing a clean, unfragmented, high-molecular weight band. Nanodrop spectroscopy indicated the purity and concentration of the DNA. From this sample, 50 ng/µL of the highest quality sample was submitted to Plasmidsaurus (South San Francisco, CA 94080, USA) for ONT and Illumina™ paired-end short-read sequencing.

### Genome assembly

Genomic DNA was submitted to Plasmidsaurus (South San Francisco, CA, USA) for combined Oxford Nanopore Technologies (ONT) long-read and Illumina™ paired-end short-read whole-genome sequencing, followed by a hybrid long-read assembly polished with short reads. ONT sequencing yielded 131,854 reads totalling 452,059,583 bp (longest read 110,884 bp), corresponding to approximately 36× raw long-read coverage of the genome. Low-quality and short reads were removed using nanoq-v0.10.0 (12) and the filtered long reads were assembled *de novo* using Flye-v2.9.1(13) with parameters selected for high-accuracy ONT reads. Illumina paired-end sequencing were quality- and adapter-trimmed using FastP-v0.23.4 (14) (Phred quality ≥15, minimum length 50 bp, poly-G trimming), retaining 8,367,000 reads (mean length 148 bp) after filtering. Quality-filtered short reads were aligned to the draft long-read assembly using bwa-mem2 (15), and the resulting alignments were used to polish the assembly at the base level with Polypolish (16) (careful mode applied at positions with short-read coverage below 25×). Assembly graph structure and contiguity were visualised using Bandage-v0.8.1 (17). Assembly completeness was assessed using BUSCO-v5.8.0 (18, 19) in genome mode against the saccharomycetes_odb10 lineage dataset (2,137 BUSCO groups), using miniprot for eukaryote gene-completeness scoring.

### Phylogenomic analysis

Thirty-five fungal genomes were selected from GenBank to represent phylogenetically diverse taxa within the sub-phylum *Saccharomycotina*. Genome assemblies were downloaded in FASTA format and assessed for completeness using BUSCO v5 in genome mode with the saccharomycetes_odb10 dataset (2,137 BUSCOs). Two genomes (GCA_030570935.1 and GCA_001984395.2) exhibited high levels of duplicated BUSCOs (>20%) and were excluded from further analyses to minimize artefacts associated with genome duplication or assembly quality. The final dataset comprised 33 genomes.

Single-copy complete BUSCO protein sequences shared across all retained 33 genomes were extracted, yielding 920 BUSCOs shared among all taxa. Each BUSCO was aligned independently using MAFFT v 7.526 with automatic parameter selection (--auto), and alignments were trimmed using trimAl v 1.4.1r22 with the -automated1 option. The trimmed alignments were then concatenated into a supermatrix comprising 373,490 amino acid positions.

Maximum-likelihood phylogenetic inference was performed on the concatenated alignment using IQ-TREE v2.2.0.5. Analyses were conducted under the LG substitution model with empirical amino acid frequencies and a FreeRate model with eight categories (LG+R8+FO), with weighted site rate optimization enabled (-wsr). Branch support was assessed using 1,000 ultrafast bootstrap replicates. The phylogeny was rooted using *Wickerhamomyces anomalus* (GCF_001661255.1) as the outgroup. All analyses were run on a high-performance computing cluster using 32 CPU cores and a memory limit of 150 GB. Resulting trees were visualized and annotated in RStudio v 2026.01.0 (R v 4.5.0) with ggtree v 3.16.3.

### Computational annotation

The predicted proteome of *P. hangzhouana* was functionally annotated using complementary orthology-based and domain-based approaches to maximize annotation depth and reliability. Prior to functional annotation, genome and gene model files were curated and standardized using Funannotate (v1.8.17)(20). Protein sequences used for annotation were generated by translating predicted coding sequences using EMBOSS (transeq) (21); sequence feature extraction was performed with BEDTools (22).

Orthology-driven functional assignments were performed using eggNOG-mapper v2 (23), using DIAMOND for accelerated protein alignments (24). This enabled assignment of Gene Ontology (GO) terms, KEGG Orthology (KO) identifiers, and Clusters of Orthologous Groups (COG) classifications. Protein domains and functional signatures were identified using InterProScan v5 (25) with annotations interpreted in the context of the InterPro resource (26). InterPro integrates multiple member databases (e.g., Pfam, SMART, PROSITE, PRINTS, Gene3D, NCBIfam, HAMAP), providing complementary evidence to orthology-based predictions for robust functional characterization.

### Construction of vectors

Plasmids for recombinant protein expression in *P. hangzhouana* were constructed using a modular cloning system with Golden Gate assembly protocols based on Type IIS restriction enzymes(27). The EP cloning system is a modular cloning strategy that was developed for the purpose of overexpressing multiple genes simultaneously(27, 28). The Orange Fluorescent Protein (OFP) gene, from anemone *Cerianthus* sp. of interest was commercially synthesised using a *Pichia pastoris* codon bias (proprietary software) for optimised expression(29). Constructs were assembled into suitable backbone BB1-acceptor vectors (C2, C14, C15, or C16) that included standard TDNA based flanking regions using Type IIS restriction cloning with the *BSA*I enzyme for EP Golden Gate construction. The kanamycin resistance gene was included in all BB1 acceptors vectors. The plasmid ID and description used in this work is presented in Figure **x** (single gene expression vector)(30).

DNA sequences encoding OFP were linked with the *P. hangzhouana* FRR5246 strains GAP1 promoter and followed by the *S. cerevisiae* CYC1 terminator. These components were cloned into basic BB1 level backbone C15, conferring geneticin resistance. Gene sequences were codon-optimized and synthesized by Azenta Life Sciences. After assembly, the whole plasmid DNA was sequenced using amplification-free long read sequencing of a linearized plasmid library (Oxford Nanopore Technologies, Plasmidsaurus, UK). The Type II restriction enzymes *Bsa*I and *Bpi*I, which cut outside of their recognition sites, are used in Type IIS restriction enzyme-compatible cloning to facilitate smooth cloning, the assembly of numerous DNA pieces, and effective one-pot cloning operation(31). In this study, we developed a Type IIS restriction enzyme-compatible cloning system specifically adapted for application in *P. hangzhouana*, as illustrated in Figure **x**. The Type IIS restriction enzyme-compatible assembly reaction involved combining the following components: 1μL of *Bsa*I or *Bpi*I (10 U), 1 μL of T4 Ligase (40 U), 1.5 μL of BSA Buffer (10×, NEB), 1.5 μL of 10 x T4 DNA Ligase Reaction Buffer, and 100 ng/µl dilutions of each insert and recipient backbone, resulting in a total volume of 15 μL. The reaction was incubated through a series of cycles: 8 to 50 cycles, depending on the number of inserts, each comprising 2 minutes at 37 °C and 16 °C, followed by 10 minutes at 37 °C, 30 minutes at 55 °C, and a final step of 10 minutes at 80 °C for ligation, digestion, and heat inactivation. This method enabled efficient assembly of DNA fragments into the desired constructs(31).

### Transformation of *P. hangzhouana*

Electrocompetent cells were prepared following the BEDS-Cereghino method (20), and stored at-80°C. The plasmids of interest (∼10µg DNA) were then linearized at the genome integration locus using a suitable restriction enzyme and transformed into electro-competent *P. hangzhouana cells* using a standard *P. pastoris* protocol modified for *P. hangzhouana* transformation(32), with modified electroporation conditions: 1.8 kV, 200 Ohms, 25 uF capacitance, (GenePulser II, BioRad). Transformed colonies were selected on YPD agar media containing 400 µg mL^−1^ geneticin (Invitrogen, Australia).

### Strain growth in BMM with alternate carbon sources for recombinant protein expression

*P. hangzhouana* 5 mL liquid YPD cultures were grown for 48 hours at 28 °C in a shaking incubator at 200 RPM. *P. hangzhouana* YPD cultures were then centrifuged for 15 minutes at 500 *g* and the supernatant removed, *P. hangzhouana* cells were then washed once with 5 mL of BMM media without an added carbon source. Washed cells were then centrifuged for 15 minutes at 500 *g* and the supernatant removed. Pelleted cells were then resuspended to a cellular O.D^600^ of 0.1 in BMM liquid media with the addition of a single carbon source and grown in a 48 well growth block in a benchtop incubator with shaking at 200 RPM. Single carbon sources were added to the BMM media at a final concentration of 2% glucose, 2% sorbitol, 0.13 mM potassium acetate, 1.9% sucrose, 2% glycerol, 2% fructose, and 2% xylose, respectively. Cultures were fed the chosen carbon source every 24 hours, and after 72 hours, screening for OFP expression was performed. The experiment was conducted in triplicate for each individual carbon source.

### Relative fluorescence measurements/Screening for OFP expression

Relative fluorescence/OFP expression was measured after 72 hours of *P. hangzhouana* growth in BMM with a single carbon source as described above. Relative fluorescence was measured using a SpectraMax M3 v3.0.22 instrument and software. 300 µL of *P. hangzhouana* culture was added in each well of a Optiplate 96 well, white polystyrene assay microplates. Relative fluorescence was measured at an excitation 540-550, and emission 563 with six reads from the top of the plate with 5 seconds of shaking performed prior to reading the sample, PMT and Optics were set to Auto. The assay was conducted in triplicate for each sample.

### Copy number

Genomic DNA was extracted using ISOLATE II Genomic DNA kit (Bioline Pty Ltd.; Alexandria, Australia) following the standard protocol. The copies of recombinant plasmid integrated into the genome of the yeast transformants were quantified using digital PCR (dPCR) with the QIAcuity digital system, using *P. hangzhouana* PGK as a single-copy gene as a reference, similar to work previously published by our group on *Pichia pastoris* (30).

## 3. Results

### Organism Phenotype and Morphology

The CSIRO *P. hangzhouana FRR5246* strain grown on YPD solid media for 3 days at 28 °C showed colonies with a margin. The colonies were of circular form with an umbonate appearance and an undulated margin with scalloped edges (Figure **1a**). Colony color was beige to white in appearance with a granular texture towards the central region of cellular growth. The newer daughter cells gave a smoother appearance as the colony grew outwards.

**Figure 1.**
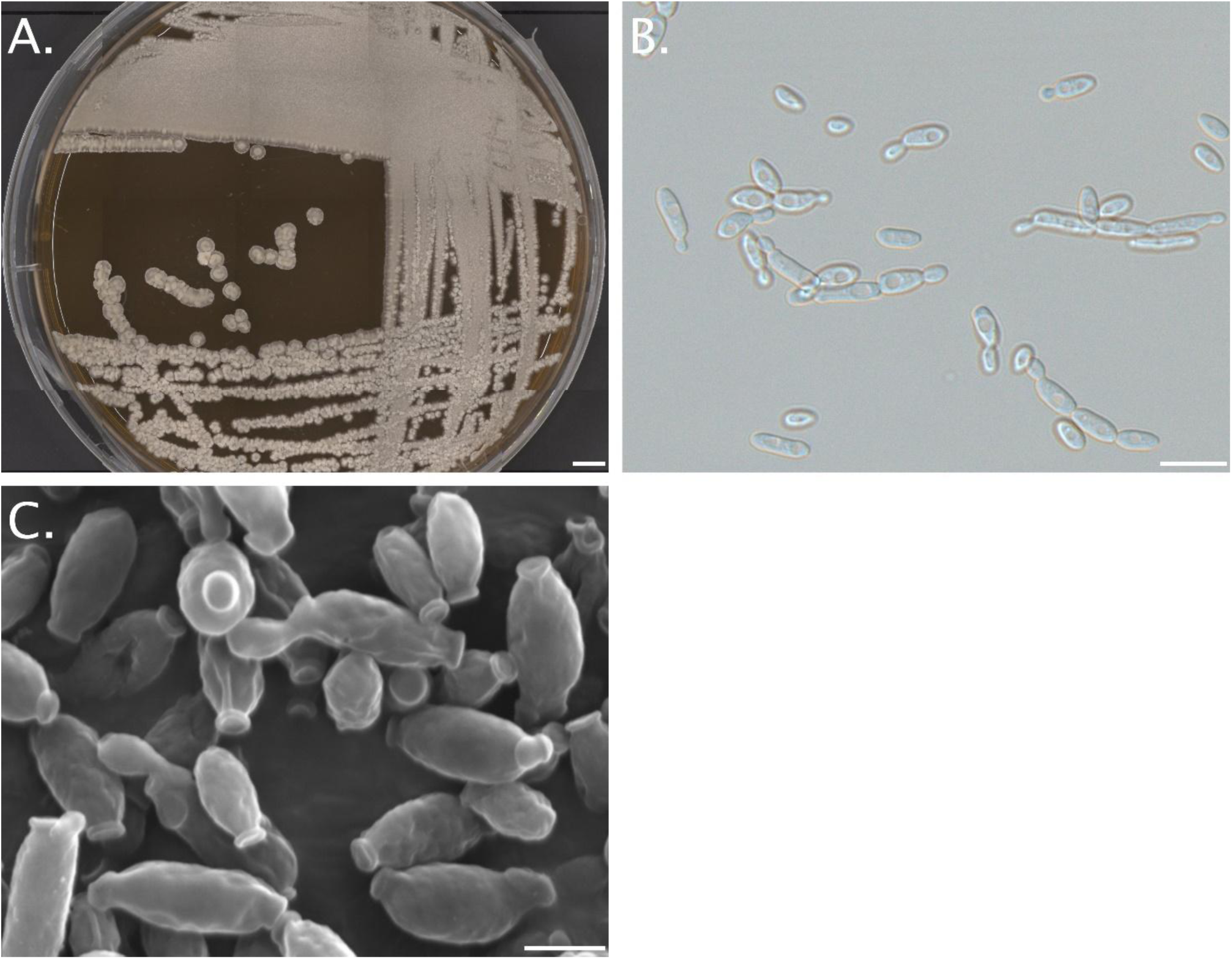
Microscopy images of P. hangzhouana. A) Light microscope image of P. hangzhouana FRR524c on YPD plate after c days of growth at 28°C, Scale bar = 5000 µm. B) DIC image of P. hangzhouana FRR524c cells after 48 hours of growth in YPD liquid culture at 28°C, with shaking at 200 rpm, Scale bar = 10 µm. C) SEM image of P. hangzhouana FRR524c cells after 48 hours of growth in YPD liquid culture at 28°C, with shaking at 200 rpm, scale bar = 2µm.

Cells grown for 3 days in liquid YPD media showed filamentous growth from budding cells, with pseudo hyphae and septate hyphae when imaged through DIC microscopy (Figure **1b**). Imaged cells were present in both an oval and elongated form as well as cells present in hyphal structures with a length of 4 to 6 µm and a width between 1 to 2 µm (Figure 1). Budding was observed in several cells, with buds emerging as small, protrusions from the parent cells indicating active cell division. These buds varied in size, indicating different stages of the budding process (Figure **1c**). The attachment points of the buds are distinct, showing a narrow neck connecting the bud to the parent cell. The cells are primarily seen in cluster/hyphal arrangements, though some isolated cells are present, suggesting localized areas of higher cell density. The *P. hangzhouana* cells in the SEM image are undergoing active budding and the attachment point of the bud to the parental cell can be clearly seen (Figure **1c**). No reproductive structures such as ascospores or asci are visible in SEM image, which is consistent with the asexual phase of growth. The cells exhibit an elongated, cylindrical morphology, which is consistent with pseudohyphal or filamentous growth of *P. hangzhouana* (Figure **1**). Individual cells measure approximately 1–2 µm in length, aligning with the expected size range for this species (Figure **1**).

### Cell Culture and Carbon Utilization

The *P. hangzhouana FRR5246* strain was grown across a range of temperatures from 25 °C to 37 °C for 48 hours. Growth was observed across all ranges tested, with the highest optical density observed when cells were grown at 25 °C (Figure **2**). Cell growth declined as the temperature increased, with the lowest optical density observed when cells were cultured at 37 °C.

**Figure 2.**
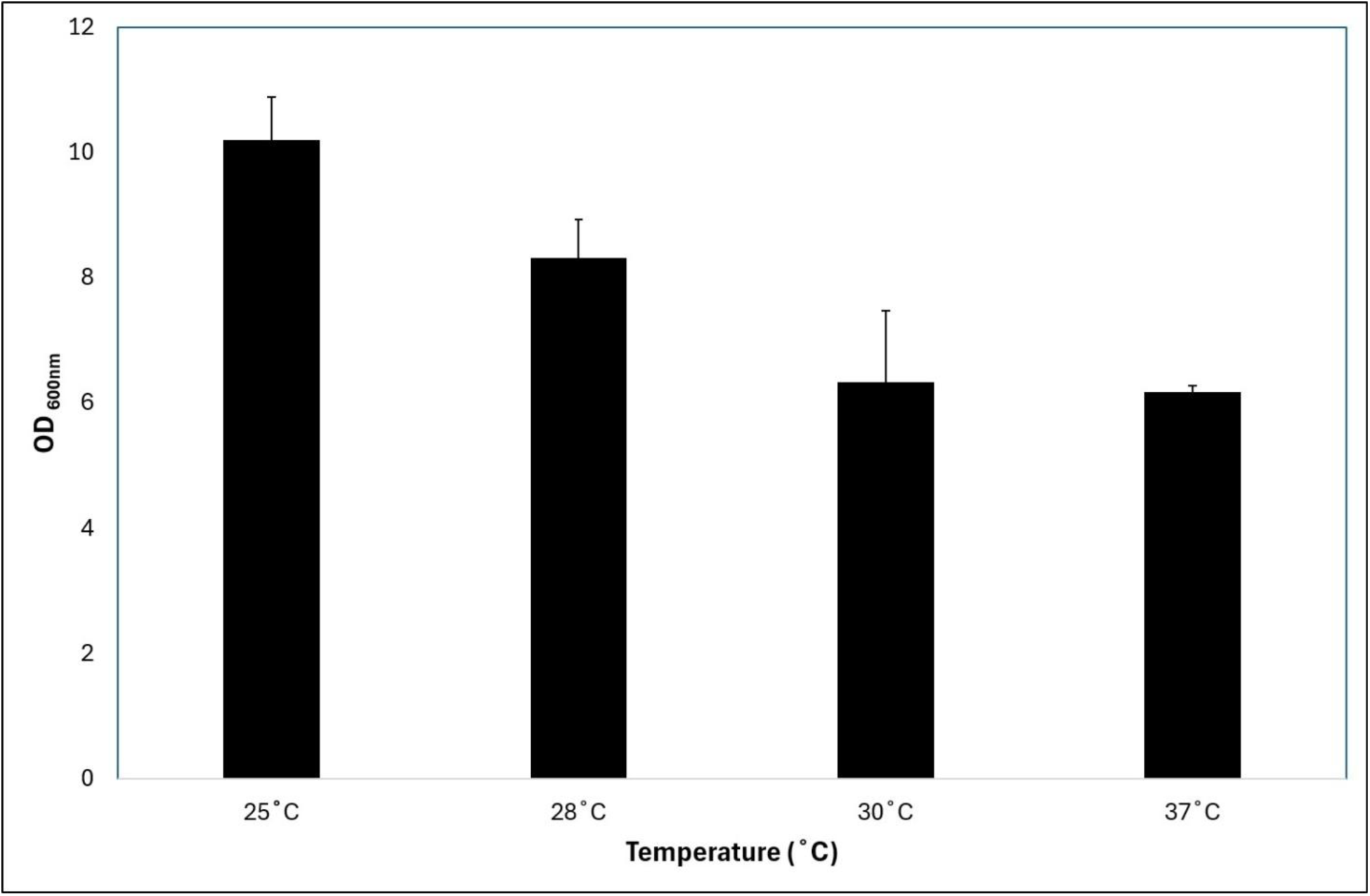
Pichia hangzhouana OD600 after 48 hours of incubation shaking at 200 rpm at temperature from 25°C to 37°C. Starting optical density OD600nm for each condition was 0.1.

In this study seven separate carbon sources were tested for the growth of *P. hangzhouana* FRR5246. The organism was grown in buffered minimal media (BM) with each separate carbon source for 48 hours starting from an OD_600_ of 0.1. The *P. hangzhouana* FRR5246 organism gave the highest optical density reading when grown in BM containing 1.9% sucrose, with an average OD_600_ reading of 32.2. The second highest OD_600_ reading was observed when cells were grown in 2% sorbitol with an average OD_600_ reading of 30.4. The third highest OD_600_ reading of 28.6 was recorded when *P. hangzhouana* cells were fed 2% fructose, followed closely by 2% glucose with a measurement of 28.4. *P. hangzhouana* cells reached an O.D^600^ reading of 24.7 after 48 hours of growth in glycerol. *P. hangzhouana* cells grew to an average OD_600_ value of 18.3 after 48 hours of growth in potassium acetate. The lowest OD_600_ values for *P. hangzhouana* cells were recorded when cells were grown in 1% methanol and when no carbon source was added to the BM media, with OD_600_ readings of 4.3 and 1.1, respectively. When fed monosaccharide and disaccharide sugars, such as fructose, glucose and sucrose, the peak cell growth or OD_600_ is reached at an earlier time point compared to the more complex carbon sources such as glycerol and sorbitol (Figure 3, B) These results indicate that *P. hangzhouana* grows well when fed both simple and complex carbon sources.

**Figure 3:**
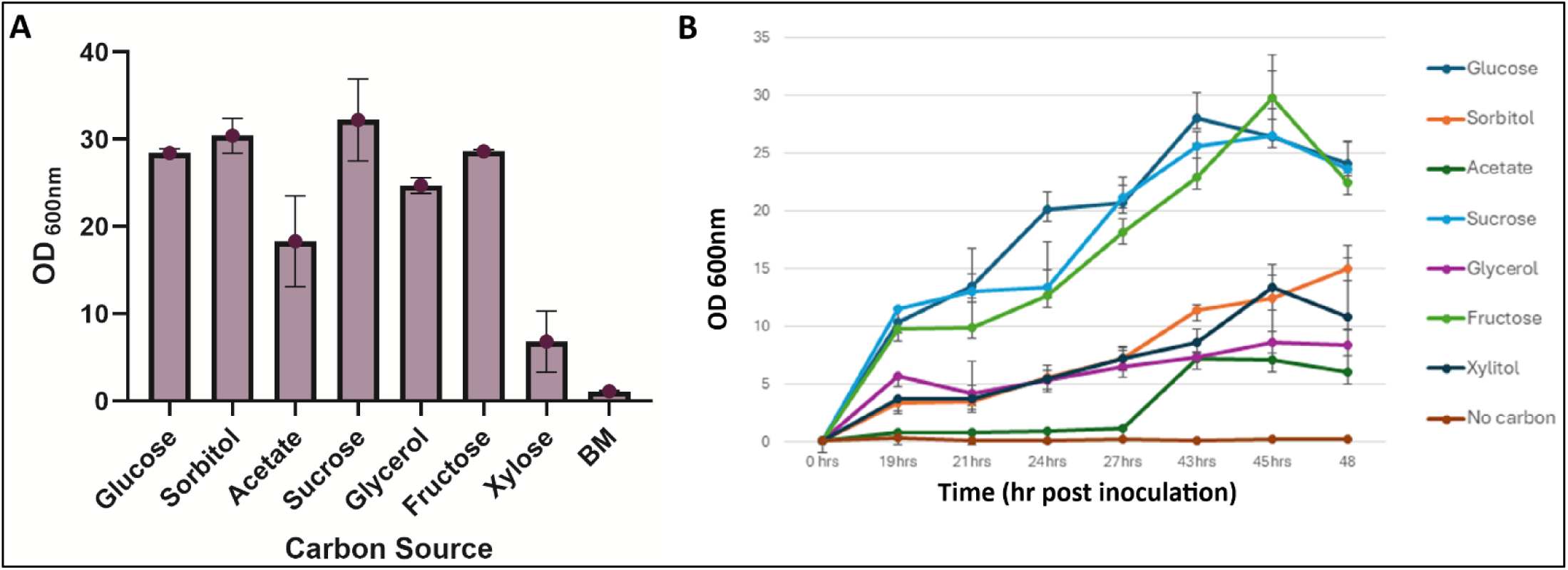
Pichia hangzhouana growth (OD600) in buffered minimal media (BM) supplemented with a single carbon source. A) P. hangzhouana growth after 72 hours in buffered minimal media supplemented with a single carbon source. B) Growth curves of P. hangzhouana cultivated in BM medium supplemented with individual carbon sources. Growth was monitored over time by measuring optical density at c00 nm reffecting biomass accumulation under each condition.

### Antibiotics susceptibility and antimicrobial activity

Antibiotics are commonly utilized in genetic engineering of yeast as a resistance cassette for selection of transformed cells. In this study the antibiotic susceptibility and resistance of *P. hangzhouana* was tested against geneticin (G418 Sulfate), hygromycin B, nourseothricin sulfate, and zeocin. *P. hangzhouana* cells were grown in liquid YPD from a starting OD_600_ of 0.1 with the addition of the chosen antibiotic, starting at a concentration of 500 µg/ml, 500 µg/ml, 250 µg/ml, and 200 µg/ml with two-fold descending concentrations for zeocin, geneticin, nourseothricin sulfate, and hygromycin B, respectively. After 48 hours of growth at 28 °C, both geneticin and nourseothricin sulfate showed inhibition of cell growth at concentrations above 31.25 µg/ml and 7.8 µg/ml, respectively (Figure 4A and Figure 4B). Zeocin concentrations above 62.5 µg/ml inhibited the growth of *P. hangzhouana* cells (Figure 4C). Hygromycin B concentrations above 25 µg/ml prevented growth of *P. hangzhouana* cell (Figure 4D). These results indicate that the *P. hangzhouana* FRR5246 strain is susceptible to hygromycin b, zeocin, geneticin, and nourseothricin sulfate at the concentrations tested in this study.

**Figure 4.**
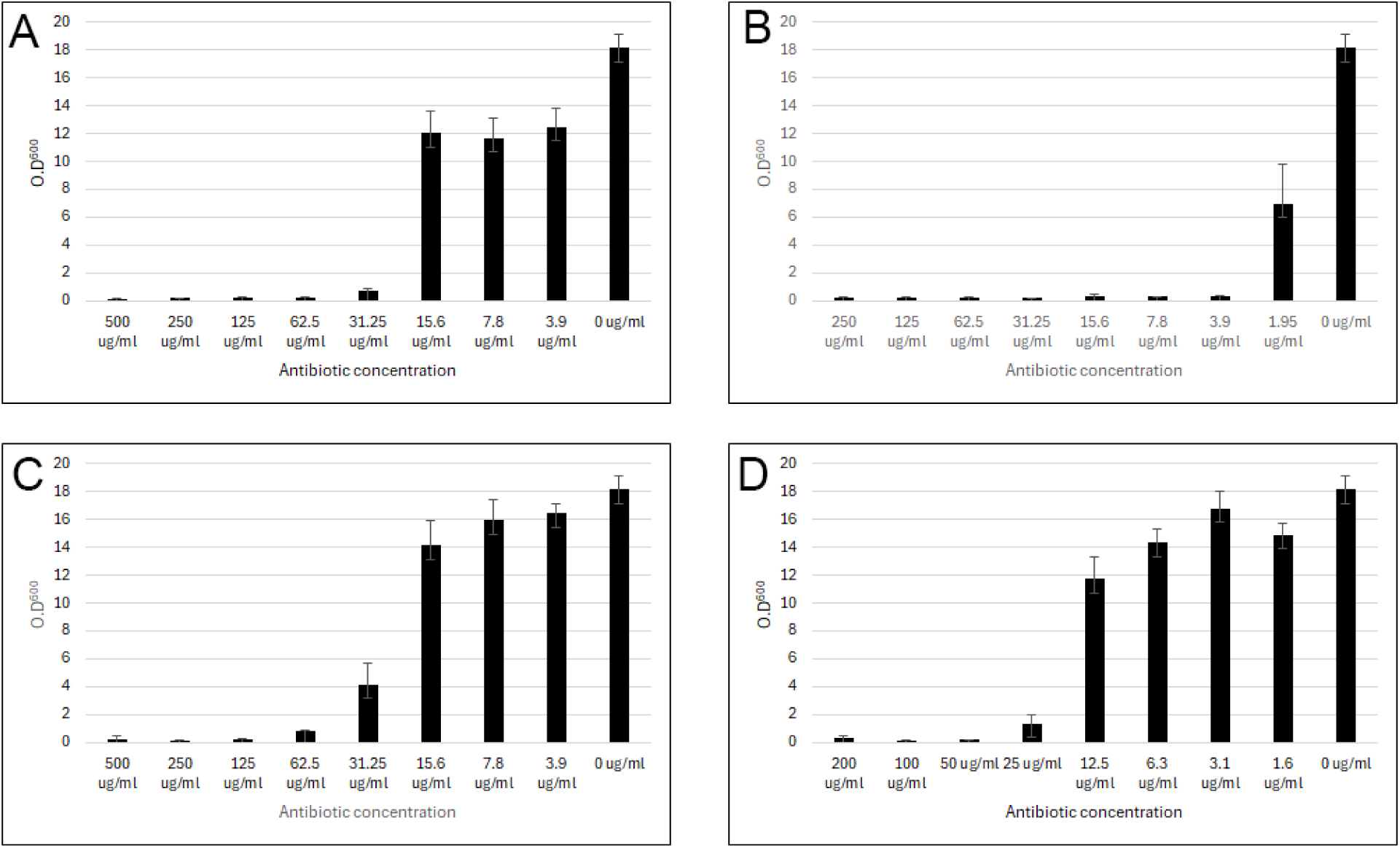
Pichia hangzhouana growth (OD_600_) in YPD media, supplement with A) geneticin, B) nourseothricin sulfate, C) zeocin, and D) hygromycin b. Cell growth measured using Optical Density^c00^ after 48 hours incubation at 28°C with shaker at 200 RPM. Concentration started at 500 µg/ml, 500 µg/ml, 250 µg/ml, and 200 µg/ml with two fold descending concentrations for zeocin, geneticin, nourseothricin sulfate, and hygromycin B, respectively.

### Antimicrobial activity

*P. hangzhouana FRR5246* cells were assessed for antimicrobial activity in a microtiter assay. In this microtiter experiment, approximately 10^5^ cells CFU/mL of *E. coli* and *S. aureus* were incubated with various concentrations of *P. hangzhouana* supernatant in Mueller Hinton Broth (MH broth) for 18 hours. The growth of bacterial cultures was first monitored by measuring the total biomass using OD measurements and confirmed by Log_10_ reduction evaluated from plate counting. The *P. hangzhouana* supernatant was tested at concentrations ranging from 180 - 1.4 mg/mL. Ciprofloxacin was used as an antibiotic control at concentrations ranging from 2 µg/mL to 0.0156 µg/mL (Figure 5). The results show that supernatant from *P. hangzhouana* has strong antimicrobial activity against *E. coli* and *S. aureus*. At the highest tested concentrations, complete growth inhibition was achieved, as indicated by 100 % inhibition based on OD measurements compared to the growth control without added *P. hangzhouana* supernatant (Table 1). Subsequent plate counting to determine the minimum bactericidal concentration confirmed a greater than 5 Log_10_ reduction in the number of viable bacteria. The significant bacterial inhibition suggests that this *Pichia* species produce highly potent antimicrobial compounds, which are effective at the concentrations tested (180 - 1.4 mg/mL).

**Figure 5.**
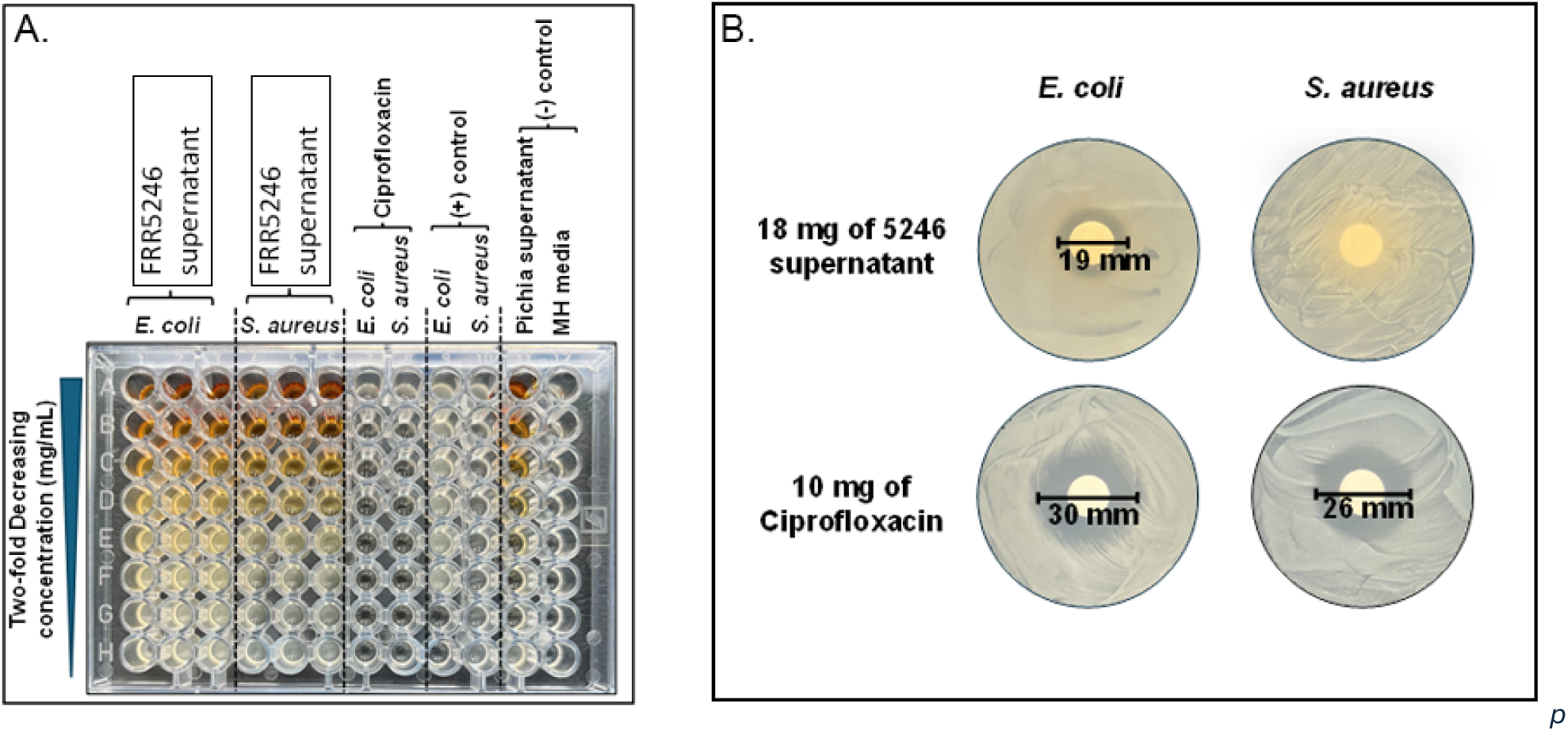
Antimicrobial activity assessment of P. hangzhouana FRR524c supernatant against both Gram-positive and Gram-negative bacteria. A) Microtiter plate-based antimicrobial assay. Two-fold decreasing concentration of test Pichia supernatants were used. B) Disk-diffusion assay on MH agar plates (150 mm diameter). For the microtiter assay Pichia supernatant tested was prepared from CSIRO FRR strain 524c at concentrations ranging from 180 - 1.4 mg/mL. Ciproffoxacin was used as an antibiotic control at concentrations ranging from 2 µg/mL to 0.015c µg/mL. 180 mg/mL of Pichia hangzhouana (CSIRO FRR 524c) was used in the Disk Diffusion assay.

**Table 1.** Growth inhibition of supernatants from both Pichia species comparing with growth control (compared to the Day 0 sample) on each strain.

| **Pichia supernatant** | **Bacteria** | **Growth inhibition calculated from OD changes comparing with growth control (%)** | **Growth/Log<sub>10</sub> reduction evaluated from plate** |
|---|---|---|---|
| 180 mg/mL of *Pichia hangzhouana* (CSIRO FRR 5246) | *Escherichia coli* | 100% | (-), > 5 Log<sub>10</sub> reduction |
|  | *Staphylococcus aureus* | 100% | (-), > 5 Log<sub>10</sub> reduction |

The highest concentration, 80 mg/mL of *P. hangzhouana*, was selected to further confirm the antimicrobial activity using a disk diffusion test (See Materials and Methods). Results of the disk diffusion test showed that *P. hangzhouana* demonstrated similar inhibition against *E. coli* as previously shown in this work (Figure 5). Although the diffusion of the *Pichia* supernatant from the disk could be seen on the *S. aureus* agar plates, no growth inhibition was apparent on this plate. This contrasts with the microtitre plate assay in broth which showed inhibition against *S. aureus*.

### Evaluation of the genome functional annotation

The predicted gene set exhibited extensive functional annotation coverage, with the majority of genes assigned to at least one conserved functional category. Comparison of Gene Ontology (GO) assignments revealed strong concordance between annotation sources, with 72.5% of genes receiving GO annotations from both EggNOG and InterPro, while smaller subsets were uniquely annotated by InterPro (14.2%) or EggNOG (6.5%). Only 6.9% of predicted genes lacked GO annotations, indicating that most predicted genes could be functionally characterized based on conserved protein domains and orthology relationships. Functional classification based on Clusters of Orthologous Groups (COG) demonstrated a broad distribution of genes across diverse biological functions (Figure 6A). The largest functional categories comprised post-translational modification, protein turnover and chaperones, intracellular trafficking, secretion and vesicular transport, translation, ribosomal structure and biogenesis, and transcription, indicating a strong representation of protein processing, intracellular transport, and gene expression functions. Genes involved in amino acid transport and metabolism, carbohydrate transport and metabolism, and energy production and conversion were also well represented, reflecting the metabolic versatility characteristic of fungal genomes. Notably, the Function unknown (S) category accounted for a substantial proportion of annotated genes (Figure 6A), representing proteins for which no reliable functional prediction is currently available. In fungal genomes, this category frequently includes lineage-specific or poorly characterized proteins that may contribute to species-specific biology and therefore represent promising candidates for future functional studies. Gene Ontology analysis further supported these findings, with the most abundant GO annotations corresponding to broad cellular component and biological process categories, particularly those associated with cellular organization and metabolism (Figure 6B). Together, the COG and GO classifications indicate that the predicted proteome is enriched in genes involved in essential cellular functions, intracellular organization, and core metabolic processes, as well as a wide range of gene-proteins involved in nitrogen metabolism.

**Figure 6.**
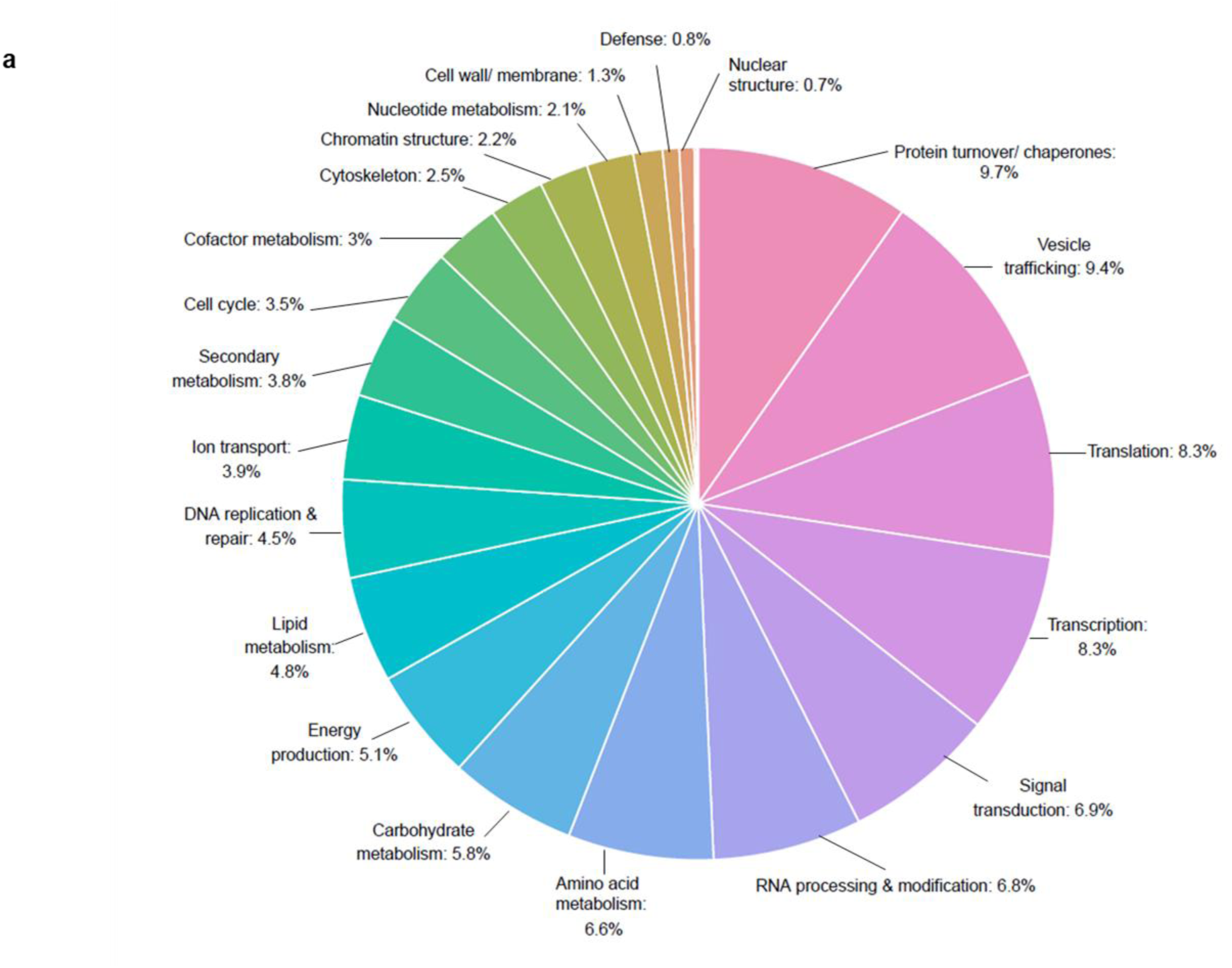
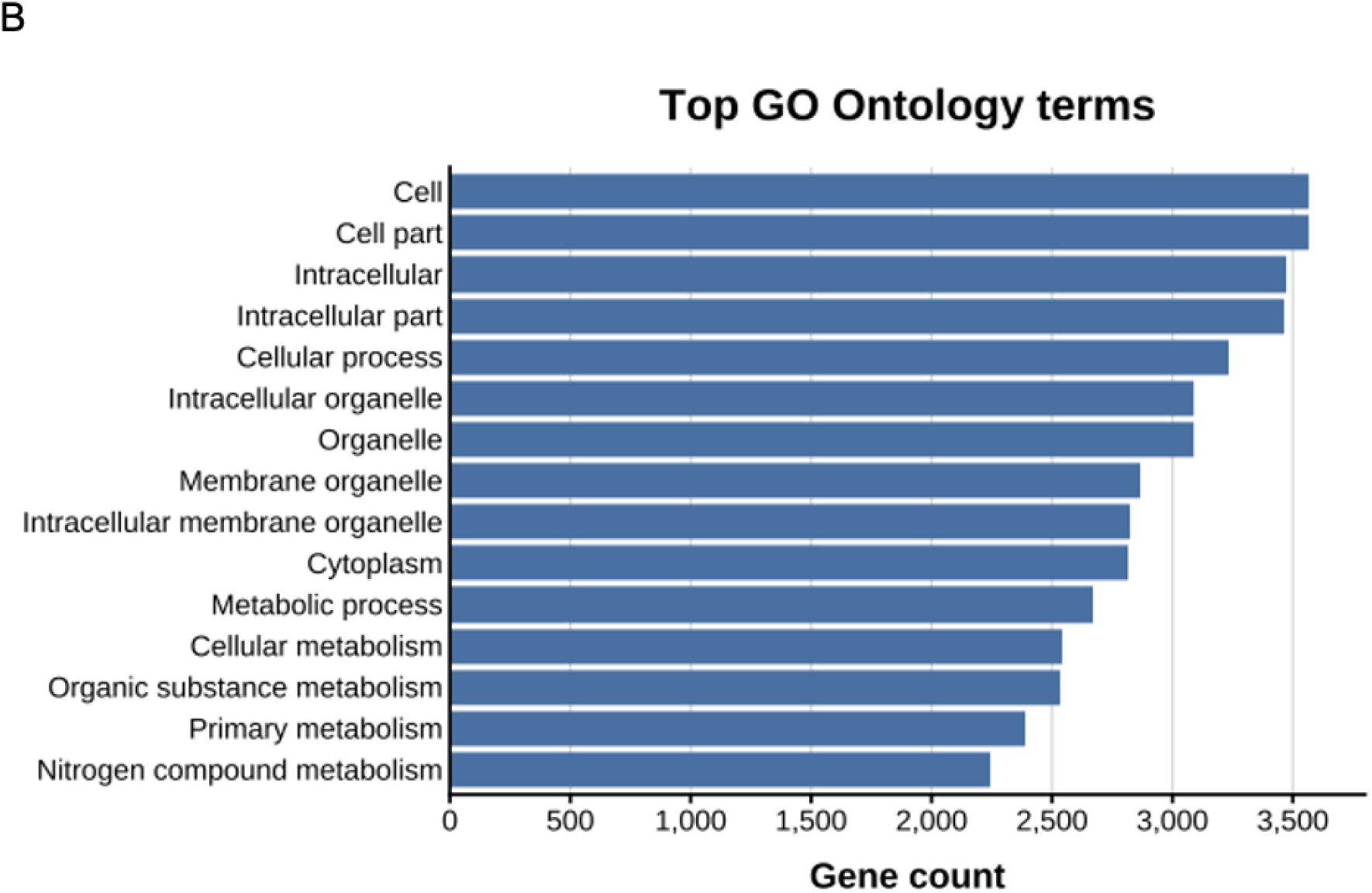
Functional landscape of the predicted proteome. (A) Distribution of predicted protein-coding genes across Clusters of Orthologous Groups (COG) functional categories. Percentages indicate the proportion of genes assigned to each functional class. (B) Most frequently assigned Gene Ontology (GO) terms among annotated genes, representing broad cellular component and biological process categories in the predicted proteome. The GO terms include cell (GO:0005623), cell part (GO:0044464), intracellular (GO:0005622), intracellular part (GO:0044424), cellular process (GO:0009987), intracellular organelle (GO:0043229), organelle (GO:0043226), membrane-bounded organelle (GO:0043227), intracellular membrane bounded organelle (GO:0043231), cytoplasm (GO:0005737), metabolic process (GO:0008152), cellular metabolic process (GO:0044237), organic substance metabolic process (GO:0071704), primary metabolic process (GO:0044238), and nitrogen compound metabolic process (GO:0006807).

### Genome assembly

The hybrid Nanopore–Illumina assembly of *P. hangzhouana* FRR5246 comprised 10 contigs totalling 12,399,881 bp (12.4 Mb), with a contig N50 of 3,746,639 bp and no internal sequence gaps. Five contigs (786,862–3,817,660 bp; 43.6–44.1% GC) accounted for the bulk of the genome and are considered nuclear in origin. Contig_1 had a distinct AT-rich composition (21.4% GC), encoding the mitochondrially-encoded gene *COX2*, and is consistent with the *P. hangzhouana* mitochondrial genome. Assembly completeness assessed against the saccharomycetes_odb10 BUSCO dataset (n = 2,137) was 87.2% complete (86.8% single-copy, 0.4% duplicated), with 0.8% fragmented and 12.0% missing BUSCOs. This genome assembly has been submitted to GenBank under BioProject PRJNA1517813 (submission SUB16433951) and is also available via FigShare (https://doi.org/10.6084/m9.figshare.33327705).

### Phylogenetic tree

To determine the evolutionary relationship of the CSIRO FRR5246 *P. hangzhouana* strain, we conducted a whole genome phylogenetic analysis. A Maximum-likelihood phylogenetic inference was performed on the *P. hangzhouana* BUSCO sequences as well as the sequences extracted from the diverse 33 genomes that were used in the analysis (See Material and Methods). The resulting tree was visualised as a rooted phylogram, with branch lengths proportional to inferred sequence divergence (Figure 7).

**Figure 7.**
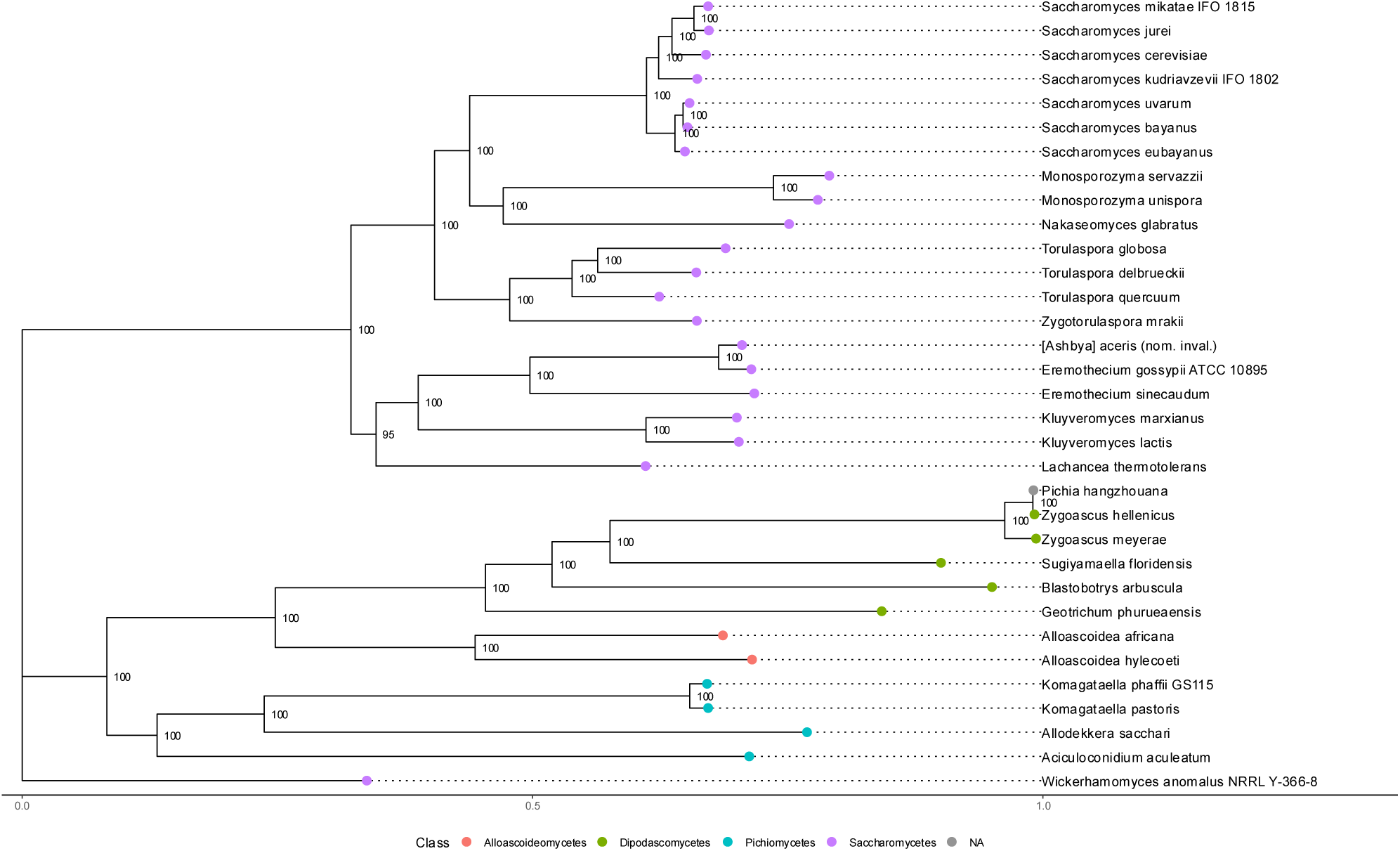
Maximum-likelihood phylogeny of Pichia hangzhouana (CSIRO FRR524c) and related yeasts based on a concatenated BUSCO protein alignment (920 genes; 373,490 amino acid positions). The tree was inferred using IǪ-TREE (LG+R8+FO; 1,000 bootstrap replicates) and rooted with Wickerhamomyces anomalus. Branch lengths indicate sequence divergence, and node labels show bootstrap support. Taxa are coloured by class, with P. hangzhouana clustering within Dipodascomycetes near Zygoascus species.

The phylogenetic analysis shows that *P. hangzhouana* clusters within *Dipodascomycetes* and is more closely related to this lineage than to *Saccharomycetes*, while *Alloascoideomycetes* form a more distantly related basal group. The *Dipodascomycetes* yeast linage represents a distinct lineage within the subphylum *Saccharomycotina*. The longer branches separating *P. hangzhouana* from *Saccharomycetes* indicate substantial evolutionary divergence. Our results presented in our analysis agree with previous reports of *P. hangzhouana* being synonymous or most closely related to the *Zygoascus species, specifically to Z. hellenicus. The next closest species* is *Zygoascus meyerae, with other members of the Dipodascomycetes clustering together, although at a further distance*.

### Secretome Analysis of *P. hangzhouana FRR5246*

Proteomic analysis of the culture supernatants from *P. hangzhouana* grown in YPD medium for 24 and 48 hours identified a total of 33 proteins. Of these, 28 proteins were detected at both time points and were considered as the core secretome of *P. hangzhouana*.

Gene Ontology (GO)-based functional classification of the 28 core secretome proteins indicated that the most represented Biological Process (BP) term was cell wall organization and carbohydrate metabolic process, each associated with ten unique proteins (Fig 8). Additional cell-wall-related terms included cell wall organization or biogenesis and fungal-type cell wall organization or biogenesis, each associated with eight proteins (Fig 8). Molecular Function annotations were mainly associated with glycoside hydrolase-related activities. After excluding broad parent GO terms, the most frequently represented terms were hydrolase activity hydrolyzing O-glycosyl compounds and hydrolase activity acting on glycosyl bonds, each associated with six unique proteins, followed by glucosidase activity, associated with five proteins (Fig 8). These results suggest that the core secretome of *P. hangzhouana* is functionally diverse, with prominent associations with carbohydrate processing, fungal cell-wall organisation/remodeling and glycoside hydrolase-related activities under the tested conditions.

**Figure 8:**
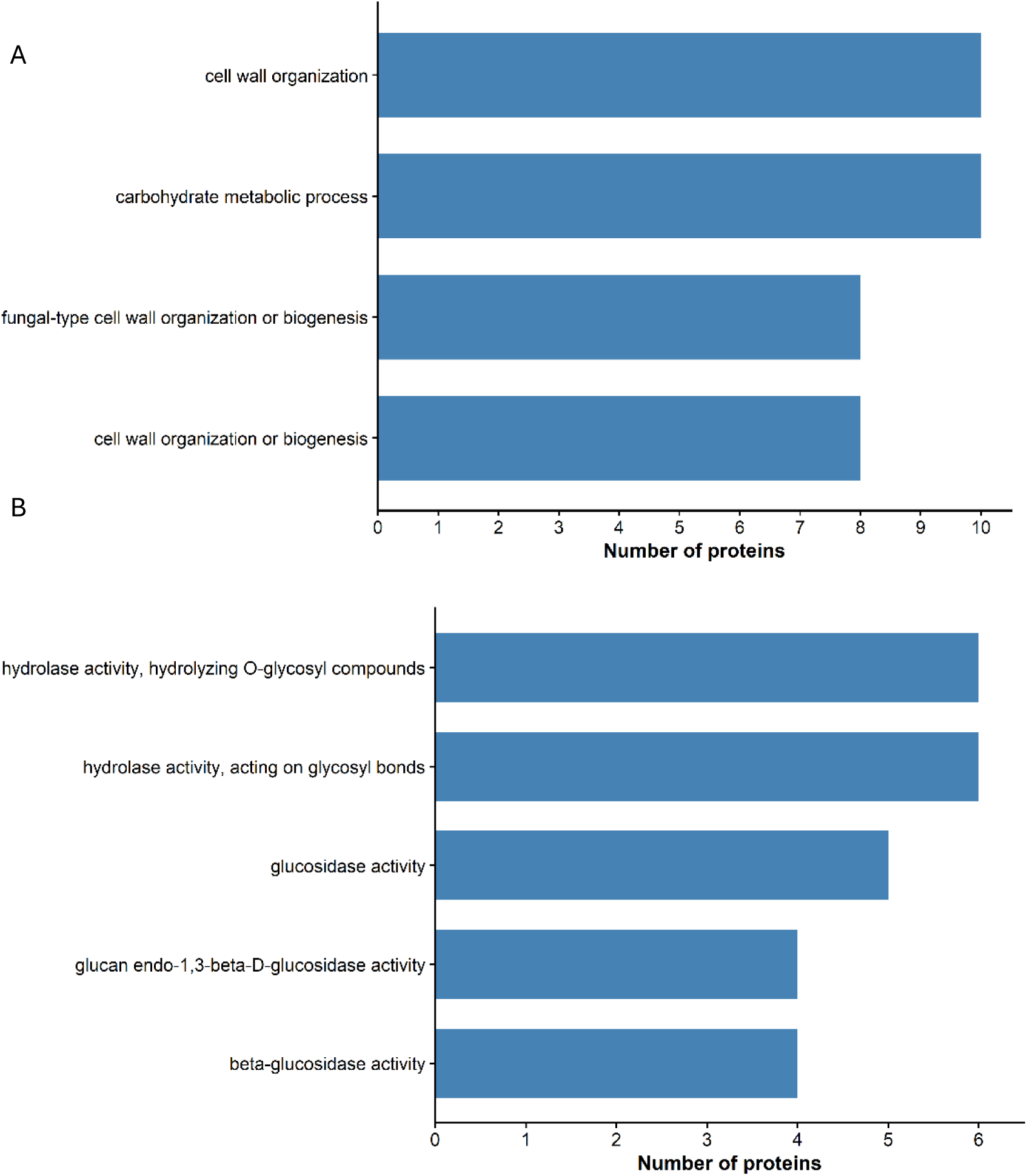
Top Gene Ontology (GO) terms associated with the core secretome of Pichia hangzhouana. The figure shows the most represented Biological Process (BP) (A) and Molecular Function (MF) (B) terms among the 28 proteins detected at both 24 h and 48 h. Bar length indicates the number of unique proteins associated with each GO term

Domain-based annotation further supported the GO-based classification. Several core Secretome proteins contained predicted carbohydrate-active and cell-wall-associated signatures including glycosyl hydrolase family 17, glycosyl hydrolase family 5, β-glucosidase-like, transglycosidase, and 1,3(4)-β-glucanase-related annotations. These predicted signatures are consistent with potential roles in carbohydrate hydrolysis, glucan turn over and cell-wall associated processes. Additional predicted domains suggested possible roles in nucleic acid degradation, phosphate metabolism, and cell envelope remodeling. Proteins with putative roles in fungal cell wall dynamics were also identified (Supplementary Table 1).

The prediction of signal peptides using the bioinformatics tool SignalP 6.0 indicated that 18 out of 28 core secretome proteins contained predicted classical N-terminal signal peptides, indicating that nearly half of the secretome may be secreted through the classical ER-Golgi-dependent pathway. supporting potential secretion through the classical ER–Golgi-dependent pathway. The remaining ten proteins did not contain predicted classical signal peptides (Supplement Table 1).

Descriptive log_2_ fold change analysis was used to compare the relative abundance of core secretome proteins detected at both 24 h and 48 h. Among the ten proteins with the largest fold changes, four showed higher relative abundance at 48 h (Fig 9). These included proteins annotated with a Kre9/KNH-like N-terminal Ig-like domain, a predicted disordered region, a Ribonuclease T2 family domain, and a dual domain profile comprising a protein kinase domain and an ARF GTPase-activating protein domain. Three of these four proteins contained predicted N-terminal signal peptides, supporting potential secretion through the classical ER–Golgi-dependent pathway (Supplement Table 1). In contrast, six proteins showed higher relative abundance at 24 h (Fig 9). These included proteins annotated with an actin-like ATPase domain, translation elongation factor 2/GTP-binding elongation factor signature, ATP synthase β-subunit/F1-ATPase β domain, and three proteins with limited functional annotation. Only one of these six proteins contained a predicted N-terminal signal peptide (Supplement Table 1).

**Figure 9:**
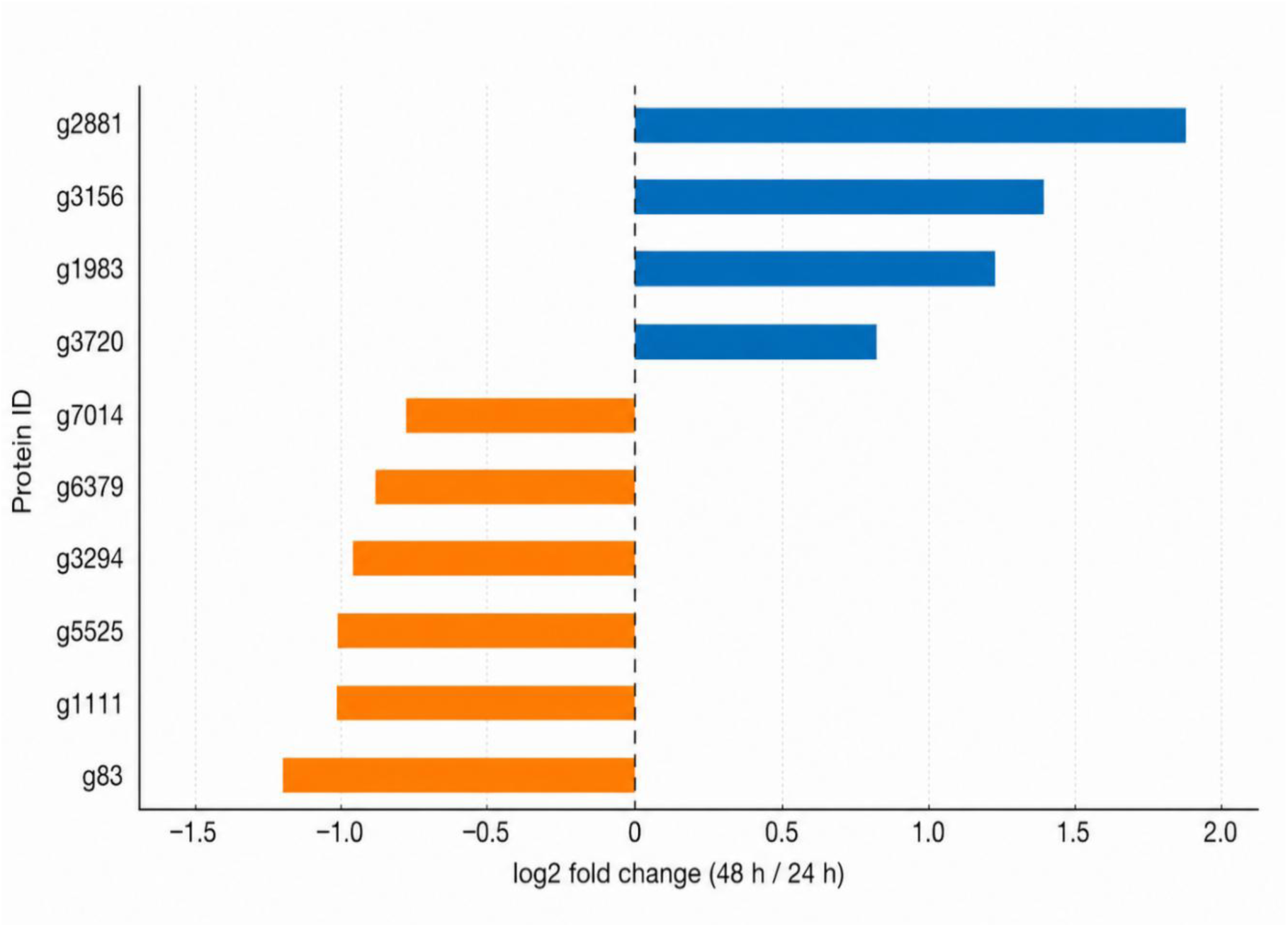
Top 10 secreted proteins showing the largest abundance shifts between 24 h and 48 h of growth. The plot shows log₂ fold changes in protein abundance (48 h vs. 24 h), based on average normalized abundances across replicates. A positive fold change (blue) indicates higher abundance at 48 hours, while a negative value (orange) reffects higher abundance at 24 hours. Predicted domain/signature descriptions and SignalP 6.1 predictions for each protein ID are provided in Supplementary Table 1.

**Figure 10.**
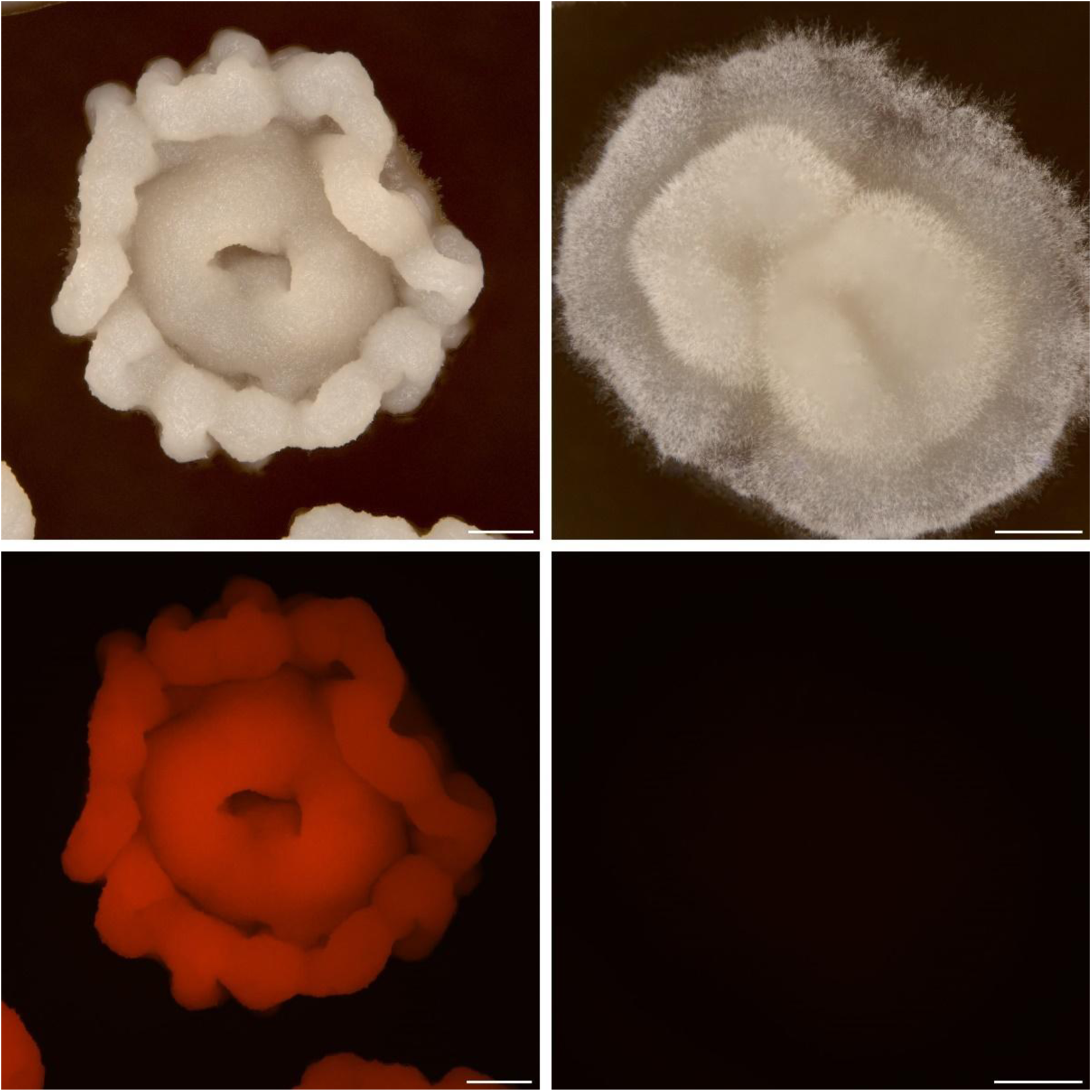
White light and Fluorescence macro imaging of P. hangzhouana wild type FRR5246 and P. hangzhouana PEP72 strains. A) White light imaging of PEP72 yeast colony on YPD media. B) White light imaging of Wild Type P. hangzhouana yeast colony on YPD media. C) Fluorescence light imaging of PEP72 P. hangzhouana yeast colony on YPD media. D) Fluorescence light imaging of Wild Type P. hangzhouana yeast colony on YPD media. Images are extended depth of focus captures. All scale bars show 500 µm

### Transformation and gene copy number

To evaluate genetic transformation in *P. hangzhouana* FRR5246, a fungal expression construct carrying an orange fluorescent protein (OFP) reporter gene under the control of the native GAP1 promoter was introduced into the strain (see Materials and Methods). Transformants were selected on YPD supplemented with 400 µg mL⁻¹ geneticin. Positive transformants exhibited cellular fluorescence, confirming successful expression of OFP (Figure X). Transformation efficiency was relatively low, with approximately 26 transformants obtained per microgram of DNA. Digital PCR analysis of a representative transformant, using the single-copy *PGK1* gene as a reference, indicated the presence of multiple integrated copies of the OFP expression cassette (33) .

The genetically modified *P. hangzhouana* colonies shown in this work expressing OFP, display notable differences in colony morphology compared to their wild type counterparts (Figure 7A and B). The GM colonies maintain a similar circular shape but consistently exhibit a more defined and wrinkled to membranous margin (Figure 7A), likely due to the expression of the recombinant protein and the associated metabolic flux changes. The presence of OFP also imparted fluorescence under appropriate lighting conditions, comparatively no fluorescence was observed in wild type *P. hangzhouana* FRR5246 colonies (Figure 7C and D).

### Expression of OFP and carbon source utilization

To investigate the effect of carbon source on recombinant protein production, the OFP-expressing *P. hangzhouana* strain (PEP72) was cultivated in BM medium supplemented with glucose, sorbitol, potassium acetate, sucrose, glycerol, fructose, or xylose as the sole carbon source. After 72 h of growth at 28°C, biomass accumulation (OD₆₀₀) and OFP fluorescence were measured from three biological replicates (Figure 11).

**Figure 11.**
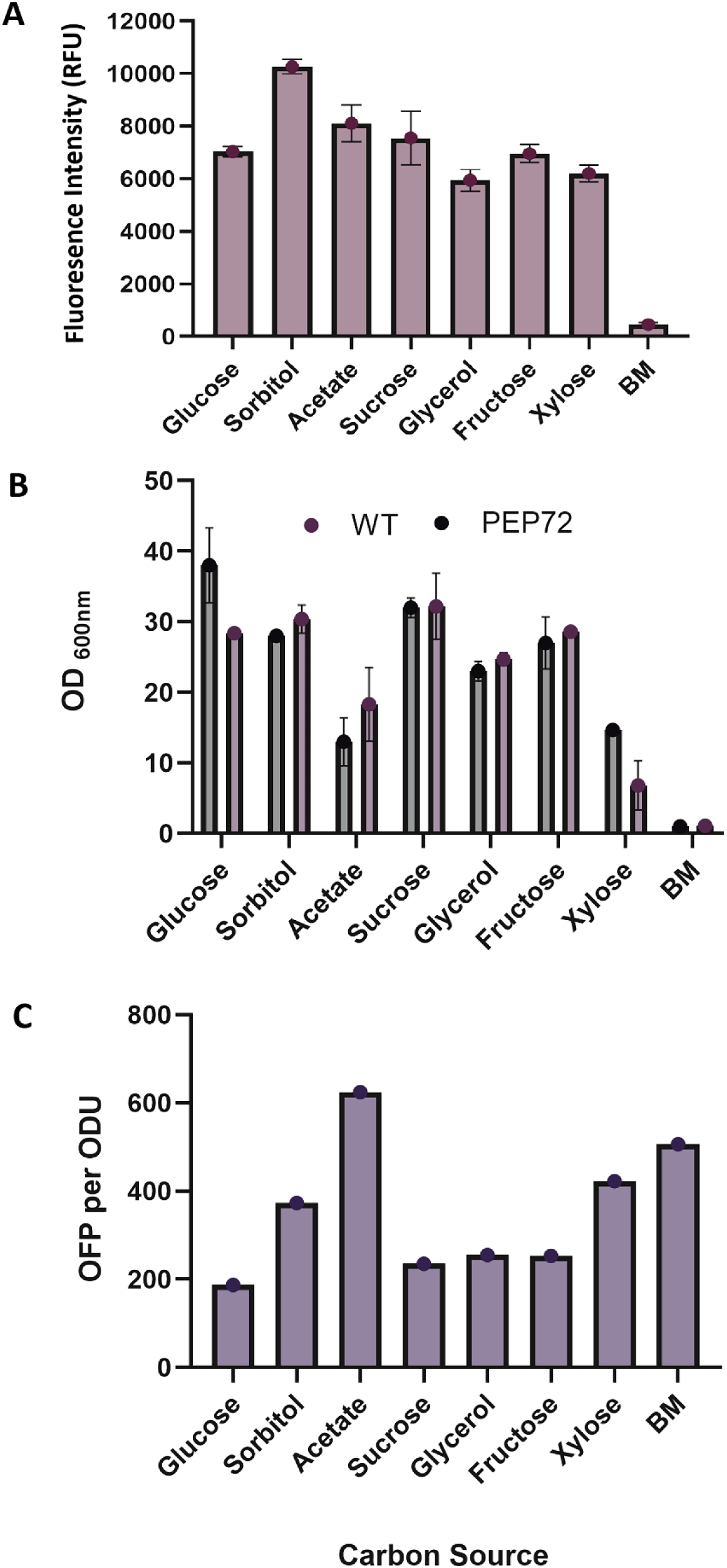
Measurement of Pichia hangzhouana PEP72 OFP expression and OD_600_ compared to the wild type strain in Buffered Minimal Media with different supplemented carbon sources. A. Pichia hangzhouana PEP72 OFP expression (RFU) . B. Comparison of OD_600_ for Pichia hangzhouana wild type and PEP72 source. C. Pichia hangzhouana PEP72 OFP expression in RFU divided by the optical OD_600_. Each Pichia hangzhouana culture was started at an optical density^600^ of 0.1 and cultured for 72 hours at 28 degrees Celsius shaking at 250 RPM.

Substantial differences in both growth and fluorescence were observed among the carbon sources tested. The highest biomass accumulation was observed in glucose-containing medium (OD₆₀₀ = 37.7), followed by sucrose (OD₆₀₀ = 32.1), whereas growth was lowest in BM medium lacking an added carbon source and was substantially reduced on acetate (Figure 11B). OFP fluorescence was detected under all growth conditions tested, confirming successful recombinant expression of the reporter gene. The highest fluorescence signal was observed in sorbitol-grown cultures (10,265 RFU), followed by acetate (8,111 RFU) and sucrose (7,551 RFU) (Figure 11A). In contrast, cultures grown in BM medium without an added carbon source produced only background fluorescence levels. No fluorescence was detected in the wild-type control strain.

To account for differences in biomass accumulation, fluorescence values were normalized to OD₆₀₀ (Figure 11C). When normalized to cell density, acetate-supported cultures exhibited the highest specific fluorescence (624 RFU OD₆₀₀⁻¹), followed by BM medium (506 RFU OD₆₀₀⁻¹), xylose (approximately 420 RFU OD₆₀₀⁻¹), and sorbitol (373 RFU OD₆₀₀⁻¹). Glucose-grown cultures showed the lowest fluorescence per unit biomass (187 RFU OD₆₀₀⁻¹). These results indicate that carbon source strongly influences recombinant protein expression in *P. hangzhouana*, with sorbitol supporting the highest overall fluorescence and acetate supporting the greatest fluorescence relative to biomass.

Comparison of the OFP-expressing strain with the wild-type strain revealed similar growth patterns across all carbon sources tested, indicating that expression of the recombinant OFP cassette did not substantially affect cellular growth under the conditions examined (Figure 11B).

## 4. Discussion

This study establishes *Pichia hangzhouana* FRR5246 as a promising non-conventional yeast for precision fermentation through the integration of genome assembly, physiological characterization, secretome analysis and genetic transformation. While *P. hangzhouana* has previously been isolated from a range of environmental and food-associated niches, its potential as a biotechnology platform has remained largely unexplored (8). Here, we provide foundational resources and proof-of-concept data demonstrating that this species can be genetically manipulated and engineered for recombinant protein production, expanding the diversity of yeast hosts available for industrial biotechnology.

A key feature of *P. hangzhouana* identified in this study is the capacity for broad carbon utilization including simple sugars (glucose, fructose), disaccharides (sucrose), sugar alcohols (sorbitol, glycerol), monosaccharides (xylose), and organic acids (potassium acetate). The highest optical density was observed when the yeast was cultured using sucrose or sorbitol as sole carbon sources, indicating efficient utilization of these carbon sources. This metabolic flexibility is consistent with previous reports describing the ability of *Zygoascus/Pichia hangzhouana* species to utilise diverse carbon substrates (34) and supports fermentation utilising a wide variety of feedstocks. For industrial biotechnology, knowledge of carbon source utilisation preferences allows for tailored fermentation strategies and provides potential opportunities to utilise renewable or waste-derived carbon sources for enhanced sustainability (10). In laboratory or industrial settings, selecting the correct carbon source can maximize growth rate and biomass yield, enhance product formation of recombinant proteins, enzymes, or metabolites and further reduce costs by enabling the use of waste-derived substrates (e.g., molasses, glycerol, acetate). *P. hangzhouana* FRR5246 grew well across all temperature conditions tested (25–37°C), demonstrating tolerance to a wide range of cultivation conditions and supporting sufficient physiological robustness for industrial fermentation processes.

*P. hangzhouana* did not utilize methanol effectively, as evidenced by its poor growth on media containing methanol as a sole carbon source. This was expected due to the absence of a methanol utilization pathway (MUT) within the organism’s genome (Genbank SUB16433951). In contrast to *P.pastoris,* which is frequently cultivated using glycerol and methanol-based processes, *P. hangzhouana* exhibited particularly strong growth on sucrose and sorbitol. Notably, both *P. pastoris and P. hangzhouana* exhibit a Crabtree-negative phenotype, meaning they do not produce ethanol under aerobic conditions, which is advantageous for high-density fermentation processes (10).

Functional genome annotation of *P. hangzhouana* FRR5246 provided a comprehensive map of its genetic capabilities, highlighting pathways involved in carbon utilization, stress response, and protein secretion. This information is critical for optimizing the yeast’s performance for precision fermentation processes and for engineering strains with enhanced traits (35). Annotation of the genome also facilitated identification of the native GAP1 promoter, which was subsequently utilised to drive heterologous expression of the recombinant OFP reporter gene.

A major outcome of this work was the successful genetic transformation of *P. hangzhouana*, which to our knowledge has not been previously reported. The antibiotic susceptibility profiling undertaken in this study identified several effective antibiotics, which could be used to screen genetic transformants for selectable resistance markers, including geneticin, hygromycin B, nourseothricin and zeocin. Utilising these findings, we demonstrated stable integration and expression of an OFP reporter construct under the control of the native GAP1 promoter using heterologous expression of geneticin antibiotic resistance as a selectable marker. Although transformation efficiencies remain relatively low compared with established yeast hosts, successful generation of fluorescent transformants provides clear evidence that effective heterologous gene expression is achievable in this species (32). These findings represent an important first step towards the development of a broader synthetic biology toolkit for *P. hangzhouana*. The recombinant protein expression studies revealed that carbon source selection significantly influenced recombinant OFP production. While the highest biomass accumulation was observed in cultures with glucose or sucrose as sole carbon source, the strongest fluorescence signal was detected in cells grown using sorbitol as carbon source. Interestingly, recombinant fluorescence normalised to biomass was highest when utilising potassium acetate as a carbon source, indicating that *P. hangzhouana* can effectively use this organic acid, which is considered a less conventional carbon source for yeast. This may be of use for future biotechnological applications involving valorisation pathways, as potassium acetate is a major by-product of various industries (36). Together, these observations suggest that recombinant protein productivity in *P. hangzhouana* is influenced not only by cellular growth but also by carbon-source-dependent metabolic regulation. Similar relationships between carbon metabolism and heterologous gene expression have been described in other industrial yeasts and highlight the importance of fermentation optimisation for maximising recombinant protein yields (37). Future work should investigate whether these observations extend to additional promoters and recombinant proteins and determine the underlying regulatory mechanisms responsible for these responses. The distinct morphological differences observed between wild type and GM *P. hangzhouana* colonies are unusual and underscore the impact of genetic modifications on yeast colony characteristics. Future studies could further explore the relationship between genetic modifications and colony morphology to enhance the utility of *P. hangzhouana* for various industrial applications.

The genome assembly generated in this study also provided the first opportunity to examine the genetic basis of the metabolic versatility observed in *P. hangzhouana FRR5246*. Physiological characterization demonstrated robust growth on a diverse range of carbon sources, including glucose, fructose, sucrose, sorbitol, glycerol, xylose and acetate, suggesting the presence of a broad repertoire of transporters and central carbon metabolic pathways. Consistent with this phenotype, functional genome annotation identified genes associated with carbohydrate metabolism, organic acid utilisation and sugar transport, supporting the ability of *P. hangzhouana* to exploit chemically diverse substrates. The COG and GO functional classifications further demonstrated that the predicted proteome is enriched in genes involved in intracellular organisation, protein turnover, transport, and core metabolic processes, reflecting a cellular architecture well suited to adapting to diverse nutritional environments (38, 39).

Metabolic versatility is a desirable characteristic for microbial cell factories because it enables the utilization of a broad range of inexpensive and renewable feedstocks, including agricultural residues and industrial by-products, thereby improving the economic and environmental sustainability of fermentation processes (40, 41). The ability of P. *hangzhouana* to efficiently utilize both fermentable sugars and non-fermentable carbon sources suggests a metabolic strategy more characteristic of non-conventional yeasts than the glucose-specialized model yeast *Saccharomyces cerevisiae*. Similar physiological flexibility has been reported for industrially important yeasts such as *Komagataella phaffii* (formerly *Pichia Pastoris*) and *Yarrowia lipolytica*, whose broad substrate utilization and respiratory metabolism have contributed to their successful application as hosts for recombinant protein production and metabolic engineering (42–44). Although the present analyses provide strong genomic evidence supporting the observed phenotype, the specific transporters, regulatory networks, and metabolic pathways responsible for substrate assimilation remain to be experimentally validated. Future transcriptomic, proteomic, and functional genetic studies will facilitate identification of the genes governing carbon utilization and metabolic regulation in *P. hangzhouana*. Together with the successful genetic transformation demonstrated in this study, these findings establish *P. hangzhouana* as a promising non-conventional yeast platform for precision fermentation and provide a foundation for future metabolic engineering and recombinant protein production.

The secretome of *P. hangzhouana FRR5246* strain exhibited a relatively limited protein profile, dominated by carbohydrate-active and cell-wall-associated proteins, suggesting a yeast-like extracellular profile mainly linked to carbohydrate processing and cell-wall remodelling under the tested growth conditions. Such a profile is consistent with reports for *Pichia pastoris/Komagataella phaffii*, where the endogenous secretome is relatively small and the conserved core secretome is mainly composed of cell-wall-associated proteins (45). This is also consistent with the broader view that *K. phaffii* has high secretory capacity but releases only low amounts of endogenous proteins, a feature that has contributed to its widespread use as a host for recombinant protein production(46). In *P. hangzhouana*, the detection of glycoside hydrolase-, β-glucosidase-, β-glucanase- and transglycosidase-associated signatures points to extracellular functions related to carbohydrate processing, glucan turnover and fungal cell-wall remodelling. These features also align with broader fungal secretome patterns such as those observed in filamentous fungi like *Trichoderma spp.*, whose secretomes are strongly enriched in carbohydrate-active enzymes involved in plant biomass and fungal cell-wall degradation (47, 48). However, unlike *Trichoderma spp.* the *P. hangzhouana* secretome is more limited, demonstrating a yeast-like extracellular profile associated mainly with carbohydrate processing and cell-wall remodelling under the tested YPD conditions.

The time-dependent secretome profile further suggests a shift in the nature of extracellular proteins between 24 h and 48 h. Proteins enriched at 48 h more frequently contained predicted N-terminal signal peptides, supporting their secretion through the classical ER-Golgi pathway. This later-stage increase in signal peptide-containing proteins is consistent with active secretion of proteins involved in cell-wall organisation and extracellular processing. For example, the increased abundance of a Kre9/KNH-like protein supports a potential role in β-glucan-related cell-wall structure, as KRE9/KNH proteins have been linked to fungal cell-wall structure and β-glucan biology(49). By contrast, several proteins enriched at 24 h were annotated as intracellular or housekeeping proteins and lacked predicted signal peptides. These proteins should be interpreted more cautiously, as proteins without classical signal peptides may arise from unconventional secretion, extracellular vesicles, or low-level cell lysis(50).

In addition to the proteomic findings, the cell-free culture supernatant of *P. hangzhouana* inhibited the growth of both Gram-positive and Gram-negative bacteria, with MIC and MBC assays supporting antimicrobial potency. This was supported by several COG and GO classifications within the proteome which predicted expression of defense-related proteins (0.8% proteome). These findings indicate that extracellular products released by this strain may possess bioactive antimicrobial functions with potential implications for food preservation, biocontrol and biomedical applications (51–53).

Collectively, the secretome profile and antimicrobial activity provide rationale for considering *P. hangzhouana* FRR5246 in future biotechnology and precision fermentation studies. A limited endogenous secretome may be advantageous for strain development because it reduces background protein complexity in the culture supernatant, which can simplify downstream processing and product recovery. The presence of SignalP-positive proteins further supports the existence of an active classical ER-Golgi secretory route, which is relevant for recombinant protein secretion, as signal peptides are commonly used to direct heterologous proteins into the secretory pathway in yeast production hosts such as *K. phaffii* (54). In addition, the enrichment of carbohydrate-active enzyme-associated proteins suggests extracellular functions linked to carbohydrate processing and fungal cell-wall remodelling (55). These functions may influence cell-wall architecture, stress adaptation and interactions with the extracellular fermentation environment, all of which are relevant to bioprocess performance (56). Antimicrobial activity further supports the potential for the yeast as a precision fermentation host for the production of extracellular bioactive compounds. The utilisation of *P. hangzhouana* for these applications would benefit from further evaluation under defined production conditions.

Overall, this study establishes a foundational transformation system for *P. hangzhouana*, enabling its use in synthetic biology and industrial biotechnology. Future work should focus on improving transformation efficiency, exploring inducible promoters for regulated expression, and assessing long-term stability of multicopy transformants. The development of additional genetic tools, such as CRISPR/Cas systems or inducible promoters, would further enhance the utility of *P. hangzhouana* as a chassis organism for precision fermentation.

## Acknowledgements

A CSIRO Future Protein Mission grant supported operating funding for summer studentship P.B.

University of Queensland - U.V. was supported by a University of Queensland Career Development Scholarship for collaborative activities.

---

**Supplementary Table 1:**
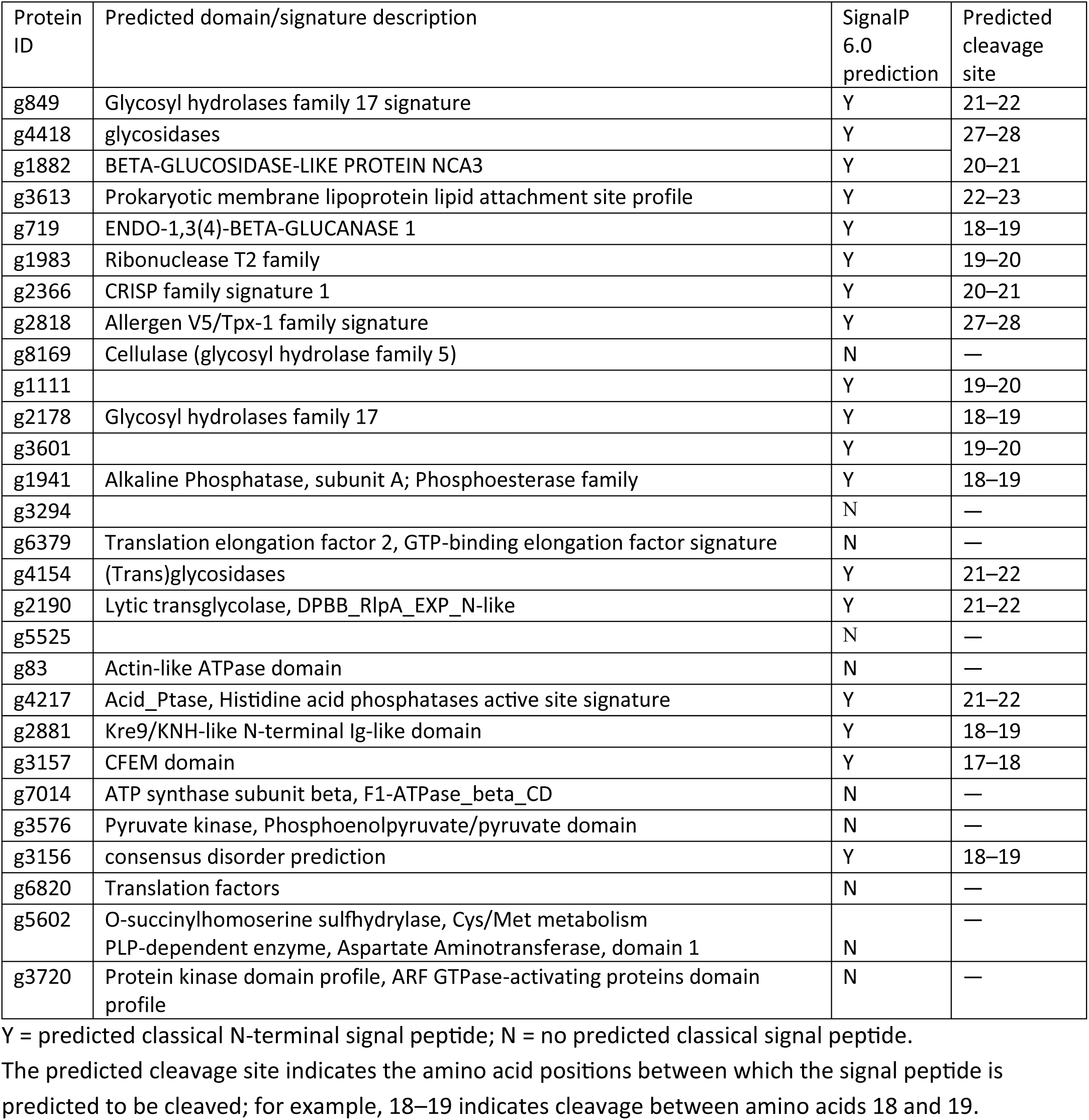
Predicted domain/signature annotation and SignalP 6.0 prediction of the 28 core secretome proteins of *Pichia hangzhouana*.

